# A ML-framework for the discovery of next-generation IBD targets using a harmonized single-cell atlas of patient tissue

**DOI:** 10.64898/2026.02.06.699999

**Authors:** Anoushka Joglekar, Ann Joseph, Pavel Honsa, Klara Ruppova, Veronica Pizzarella, Amanda Honan, Devin Mediratta, Emily Vollmer, Evan Geller, Martin Valny, Eva Macuchova, Shiwei Zheng, Alisa Greenberg, Petr Taus, Alina Kline-Schoder, Renata Konickova, Lucie Cerna, Hila Sharim, Lior Ness, Giorgio Camilli, Eleni Chouri, Irem Kaymak, Joshua D’Rozario, Daniela Castiblanco, Joao Oliveira, Francesca Prandi, Nikolay Popov, Ana Laura Moldoveanu, Christopher Oliphant, Leire Escudero-Ibarz, Florian Uhlitz, Elizaveta Freinkman, Jana Sponarova, Priyanka Vijay, Cailin Joyce, Irina Leonardi, Shikha Nayar, Tali Raveh-Sadka, Noam Solomon, Adam Platt, Tatiana Ort, Greet De Baets, Daniele Corridoni, Aleksandra Wroblewska, Adeeb Rahman

**Affiliations:** Immunai, New York, NY 10016, USA; Bioscience Immunology, Research And Development, Respiratory & Immunology (R&I), BioPharmaceuticals R&D, AstraZeneca, Cambridge, UK; Translational Science and Experimental Medicine, Research And Development, Respiratory & Immunology (R&I), BioPharmaceuticals R&D, AstraZeneca, Cambridge, UK; Bioscience Immunology, Research And Development, Respiratory & Immunology (R&I), BioPharmaceuticals R&D, AstraZeneca, Gaithersburg, US; Centre for Genomics Research, Discovery Sciences, BioPharmaceuticals R&D, AstraZeneca, Cambridge, UK

## Abstract

Target discovery for IBD has traditionally relied on genetic associations, which lack the cellular resolution needed to identify novel, actionable, cell type-specific disease pathways. Here, we describe an integrated analytical and experimental framework that leverages harmonized single-cell data to systematically discover novel therapeutic strategies for IBD.

We used AMICA DB^TM^, Immunai’s harmonized database of single-cell RNA datasets to construct a harmonized 1 million single-cell atlas of the human intestine. We applied a machine learning framework (Immune Patient Representation, IPR) to identify disease-associated transcriptional programs and cell type-specific gene targets. Candidate targets were prioritized using atlas-derived metrics, refined using custom criteria emphasizing translational actionability, and validated across independent clinical cohorts. Select candidates were evaluated in human primary-cell models reflecting the target’s cell-type context.

The IPR framework identified 85 disease-associated transcriptional programs and ranked 400 cell type-specific target genes across immune and stromal lineages. Disease-associated programs were interpreted using a structured AI-assisted reasoning framework for structured biological reasoning, linking them to IBD-relevant pathways and guiding the identification of novel, promising gene targets. Functional validation of two cell-type-specific candidates, PTGIR in myeloid cells and IL6ST in fibroblasts, confirmed the reduction of inflammatory and fibrotic pathways linked to IBD pathology. Multi-omic profiling and projection of *in vitro* phenotypes to patient datasets demonstrated the reversal of disease-associated programs via mechanisms distinct from those of existing biologics.

Our single-cell anchored, machine-learning framework integrates *in silico* discovery with experimental validation, revealing new cell type-specific therapeutic opportunities and supporting a scalable approach for precision target discovery in IBD and other immune-mediated diseases.

## Introduction

Despite recent progress in therapeutic discovery, inflammatory bowel diseases (IBD) patients still have considerable unmet clinical need, due to high rates of therapeutic non-response, relapse, and adverse events^1,2^. Identifying novel disease pathways that underlie inflammation or non-response to existing treatments is critical for the development of next-generation therapeutics. Historically, IBD target discovery has been driven by genome-wide association studies^2^ which have identified over 200 IBD-associated genetic loci^3–6^. These studies have highlighted genes involved in immune regulation, barrier function, and cytokine signaling, profoundly advancing our understanding of IBD’s genetic architecture^2–4^. However, this previous dependence on genetic targets has led to an enrichment of therapies that act on pleiotropic pathways, such as steroids, TNF blockers, and JAK inhibitors. These broad-acting therapies can cause systemic immune suppression, increased infection risk, and metabolic complications^7^. Cell type-specific therapeutics, such as vedolizumab, which selectively blocks α4β7 on gut-homing lymphocytes, are an emerging alternative class that achieve potent effects in disease-relevant cell-types, while minimizing undesired effects in other cell-types or tissues^8^.

Clinical research in IBD increasingly incorporates single-cell RNA-Sequencing (scRNA-seq). This type of data provides the resolution needed to identify disease-relevant transcriptional networks and therapeutic targets within specific immune and stromal cell subsets^9,10^. It can also be used to develop and refine preclinical models that accurately recapitulate key molecular features of disease, promoting the translatability of target-validation experiments^11^. However, the usefulness of existing scRNA-seq data is limited by the prevalence of small cohorts from different research centers, technical variability, and difficulties in dataset harmonization^9,12,13^.

To overcome these challenges, we leveraged AMICA^TM^ platform, Immunai’s single-cell data infrastructure that includes 400 proprietary and public human single-cell studies, harmonized with a pipeline that integrates and annotates single-cell datasets at scale, ensuring comparability of cell-type and metadata across studies. For this study, we extracted more than 200 intestinal samples and nearly 1 million cells from the AMICA database (AMICA DB^TM^) to generate a comprehensive single-cell intestinal atlas encompassing immune and stromal compartments.

The AMICA^TM^ platform encompasses a set of tools that model single-cell data at the cell, sample, or patient level depending on the biological question. AMICA^TM^ platform-based models are being built to learn and represent the manifold of multimodal, multi-sample patient data as the Immune Patient Representation (IPR), wherein similar immune profiles are placed in close proximity in a shared latent space. In this study, we used a transcriptomic-only version of IPR designed to work with aggregated gene expression profiles for each cell-type within a sample. We applied linear dimensionality reduction to identify robust gene programs that consistently distinguish disease phenotypes across samples in a cell-type resolved manner. These gene programs were evaluated with a structured reasoning framework that distills IPR outputs into biologically coherent processes enabling the translation of statistical signals into mechanistic hypotheses and cell-type–specific therapeutic targets.

Our analysis prioritized a set of 19 candidate gene targets with a strong weight of evidence and a strong fit to our desired target product profile. These targets underwent functional validation using disease models that were optimized using computational methods to reflect disease-relevant cellular states. Here, we present two genes, PTGIR in macrophages and IL6ST in fibroblasts, as functionally-validated cell type-specific targets poised for successful clinical development.

## Materials and Methods

### IBD Single-Cell Atlas Construction

Using Immunai’s Annotated multi-omic Immune Cell Atlas (AMICA DB^TM^), we constructed a harmonized IBD tissue atlas from 20 publicly available scRNA-seq studies. This atlas comprised 530 intestinal samples (**Table S1**). Studies were required to have UMI (Unique Molecular Identifier)-based library types, and available raw data. Briefly, data ingestion into AMICA DB^TM^ includes the processing of raw FASTQ files using 10x Genomics Cell Ranger (v5.0.1) to generate count matrices. Low-quality cells are removed using standard filters (≥200 genes, ≥300 UMIs, <10% mitochondrial UMIs), and potential doublets are excluded. The data were integrated, normalized, and batch-corrected for sample-specific effects. After integration, cluster based annotation leveraged our proprietary tissue reference and cell-type ontology. The final integrated object contained 994,206 cells spanning 129 distinct cell subtypes and states.

### Immune Patient Representation Framework

To derive robust, sample-level biological signals, we learned an Immune Patient Representation (IPR) from a subset of eight studies from the IBD Single-Cell atlas (**Table S1**). These studies were selected to ensure the availability of disease-relevant metadata and a comprehensive representation of intestinal cell types, while excluding datasets with sorted cell populations that could bias cell-type representation and introduce technical artifacts. We employed a transcriptomic-only version of IPR trained only on public intestinal single-cell transcriptomic data. IPR estimation comprised three steps: A) Pseudobulk construction: nonlinear batch correction followed by aggregation of gene expression by annotated cell type within each sample. B) Linear dimensionality reduction: to obtain latent components capturing major biological variation. C) Feature extraction: Use cell-type–specific gene loadings from these components as IPR features (gene sets) for downstream analyses. We evaluated IPR features across 12 predefined biological contrasts (**Table S2,** e.g. inflamed vs. non-inflamed, anti-TNF responder vs. non-responder) using Wilcoxon rank-sum tests, considering Bonferroni-corrected p < 0.1 as significant. For anti-TNF–related signatures, which had limited sample support, we required additional validation in external bulk transcriptomic cohorts (**Fig S3G, Table S3**).

This workflow provides a transparent, reproducible, and mechanistically interpretable instantiation of IPR for target discovery using publicly available scRNA-seq datasets.

### Target Identification and Prioritization

Each top IPR feature was distilled into a ranked list of candidate genes. Genes were included if they met one of these 2 criteria:

Pearson’s correlation between gene expression and IPR feature value ≥0.6.
Pearson’s correlation between gene expression and IPR feature value ≥0.3 and Median absolute deviation (MAD) normalization of IPR features coefficients ≥ mean + 1.5sd.

To ensure comprehensiveness, this list was further supplemented with differentially expressed genes (limma-voom, n=10 DEGs). This strategy yielded a list of 4,036 candidate genes. Candidate target genes were ranked by integrating the IBD atlas derived transcriptional evidence with a wide range of orthogonal, publicly available data across four key domains:

1. genetic and functional linkage to IBD (Open Targets, Locus2Gene, GWAS catalogs),
2. network biology (STRING protein-protein interactions, MultiNicheNet cell-cell communication),
3. druggability and therapeutic potential (DGIdb, Surfaceome, known IBD/IMD drug targets), and
4. potential safety liabilities (expression in heart/liver scRNA-seq atlases, mouse phenotype data from IMPC).

Details available in **Table S4**. Each gene was scored for these metrics, which were then aggregated via a weighted sum to produce a final ranked list of targets.

### Interpretation of IPR features

To systematically interpret IPR gene-level signals into mechanistic modules in the relevant cell type-specific context, we used AMICA-Reason^TM^, our structured, agentic reasoning framework. AMICA-Reason^TM^ integrates statistical enrichment with large language model (LLM)-based functional analysis to generate both molecular- and cellular-level interpretations. The molecular module combines GSEA outputs with LLM-assisted analysis of internal and public biomedical resources (e.g., Human Protein Atlas, PubMed) to contextualize gene-set functions. The cellular module identifies recurring immune processes across IPR features within each cell type, resolves pleiotropic signals, and groups coherent functions into higher-order cellular insights. Details of AMICA-Reason^TM^ are described in the supplementary Methods. The flexibility of AMICA-Reason^TM^ allows us to supplement curated pathway libraries (GO, KEGG, Reactome) with signatures derived from single-cell and bulk transcriptomic datasets curated within AMICA^TM^ (distinct from those used to build the intestinal single-cell atlas) and thus enhance model interpretation at multiple levels.

### Gene Knockout in Primary Human Cells

Functional validation was performed using endonuclease mediated knockout in primary human cells. CD14+ monocytes were isolated from healthy donor PBMCs and differentiated into mononuclear phagocytes (MNPs) over 5 days with M-CSF. Primary human intestinal fibroblasts were expanded in culture. Both cell types were nucleofected with pre-complexed ribonucleoproteins. Following recovery and stimulation with disease-relevant cytokines, cells and supernatants were harvested for downstream analysis. Full protocol details, including reagent concentrations and sgRNA-Sequences, are available in the Supplementary Methods.

### Functional Validation

Target knockout efficiency was confirmed at the transcript level by qPCR. Cellular phenotypes were assessed by multi-color spectral flow cytometry (Cytek Aurora), and secreted protein levels in supernatants were measured using a multiplexed analyte assay (Nomic Bio). Global transcriptional changes resulting from gene knockout were profiled by bulk RNA-seq. Libraries were prepared using the Illumina Stranded mRNA Prep Kit and sequenced on an Illumina NovaSeq system. Raw sequencing data was processed through a standardized workflow to generate gene count matrices. Differentially expressed genes between target and control knockout conditions were identified using limma-voom, and pathway enrichment was assessed using GSEA.

### Clinical projection of *in vitro* signal

To validate the translational relevance of the *in vitro* results, gene signatures derived from our *in vitro* knockout experiments were projected onto the pseudobulked IBD tissue atlas. For each candidate target, a signature comprised genes significantly upregulated/downregulated post knockout. The average signature expression was calculated in the target cell type and normalized to a control signature that was derived by randomly selecting genes that matched the expression profile of the target signature. Conversely, the original IPR feature signatures were tested for enrichment in the *in vitro* knockout data to confirm that the target perturbation had modulated the intended disease-relevant pathway. Datasets used for clinical projection are listed in **Table S1**.

### Data Availability

All clinical scRNA-seq and bulk transcriptomic datasets used in this study were obtained from publicly available repositories and are listed in Supplementary **Table S1** with their respective accession numbers.

## Results

### Construction of a harmonized IBD single-cell atlas

Single-cell data presents computational challenges due to its high dimensionality, small sample sizes, and inherent noise from sequencing and batch effects. To overcome these challenges and build a comprehensive, harmonized single-cell map of the human intestine, we leveraged AMICA DB^TM^, Immunai’s harmonized database of proprietary and public single-cell RNA-Sequencing (scRNA-seq) datasets (**Fig 1A, Table S1**). AMICA DB^TM^’s standardized ingestion and integration pipeline enables the construction of a high-quality tissue atlas. We used AMICA DB^TM^ to integrate intestinal tissue samples from 20 publicly available studies into a harmonized single-cell atlas. The resulting atlas comprised 994,206 cells from 530 intestinal tissue samples, including more than 200 Crohn’s disease (CD) ulcerative colitis (UC) patients, and healthy controls (**Fig 1B, Fig S1A,B**). Cell type annotation identified 76 cell subtypes, and 127 cell subtypes and states (**Fig 1C**), whose accuracy was validated by the expression of canonical markers (**Fig 1 D, Fig S1 C,D**). This harmonized intestinal tissue atlas served as the single-cell data foundation for target discovery, providing increased resolution of disease-relevant cell types and states compared to the original individual studies (**Fig 1E**)^14^. For independent validation, we further leveraged the constantly growing AMICA DB^TM^ to assemble an independent cohort of 163 IBD and healthy intestinal samples (520,969 cells) from four newly published studies through the same harmonization pipeline (**Table S1**).

**Figure 1:**
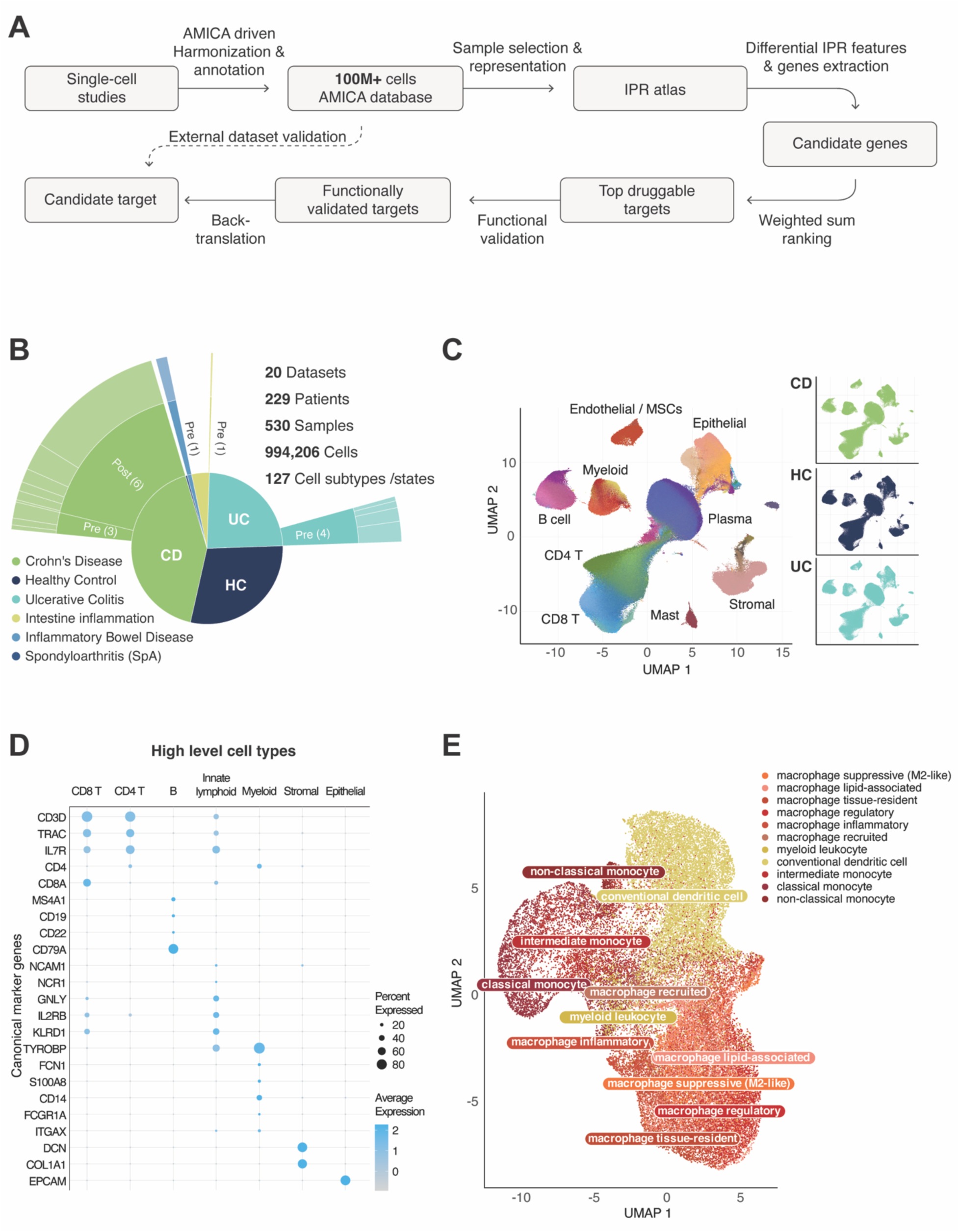
**The AMICA^TM^ Platform integrates 20 single-cell studies into a harmonized intestinal atlas** A) Overview of the AMICA^TM^-based framework used for single-cell atlas construction, IPR-driven target discovery and functional validation. B) Starburst plot summarizing sample composition across the 20 studies included in the intestinal sample atlas. Numbers in parentheses indicate the number of studies per treatment category. C) UMAP visualization of the single-cell intestinal atlas, colored by major cell type (left) and diseases category (right). D) Dot plot of canonical marker genes across major immune and stromal cell types. Dot size represents the percentage of cells expressing the gene, color indicates the pseudobulk normalized average expression level per cell type. E) UMAP visualization of myeloid cells within the single-cell IBD atlas. Cells are colored by cell type. CD, Crohn’s disease; HC, healthy control; IBD, inflammatory bowel disease; SpA, spondyloarthritis; UC, ulcerative colitis; UMAP, Uniform manifold approximation and projection; B, B cells (including plasma cells).

### Identification of cell type-specific disease pathways using an immune patient representation method

Our harmonized intestinal tissue atlas forms a scalable basis for target discovery in IBD. To extract actionable cell type-specific disease programs, we applied the Immune Patient Representation (IPR) framework, an ML framework that leverages single-cell information to learn disease-relevant transcriptional variation within and across patient samples^15^. IPR extracts this variation as ranked, cell type-specific transcriptional programs, or IPR features, which represent the strongest gene signatures associated with disease state, treatment response, or other biological conditions (**Fig 2A**). To optimize patient representation, we selected a subset of eight studies from the full intestinal atlas. This subset was chosen because it 1) provided comprehensive coverage of immune and stromal cell populations, 2) contained sufficient samples across key biological contrasts, such as inflamed versus non-inflamed states in Crohn’s disease and ulcerative colitis, 3) was derived from unsorted single-cell intestinal preparations (**Fig 2A-B, Fig S2A-C**, **Fig S3A, Table S1**). This strategy ensured robust statistical power and avoided biases from under-represented conditions and cell types while retaining the biological diversity of the full intestinal atlas. Consistent with prior literature, this subset recapitulated known IBD abundance features, such as increased infiltration of monocytes and recruited macrophages and reduced tissue-resident macrophages in inflamed samples (**Figure S2 D,E**)^16–18^.

**Figure 2:**
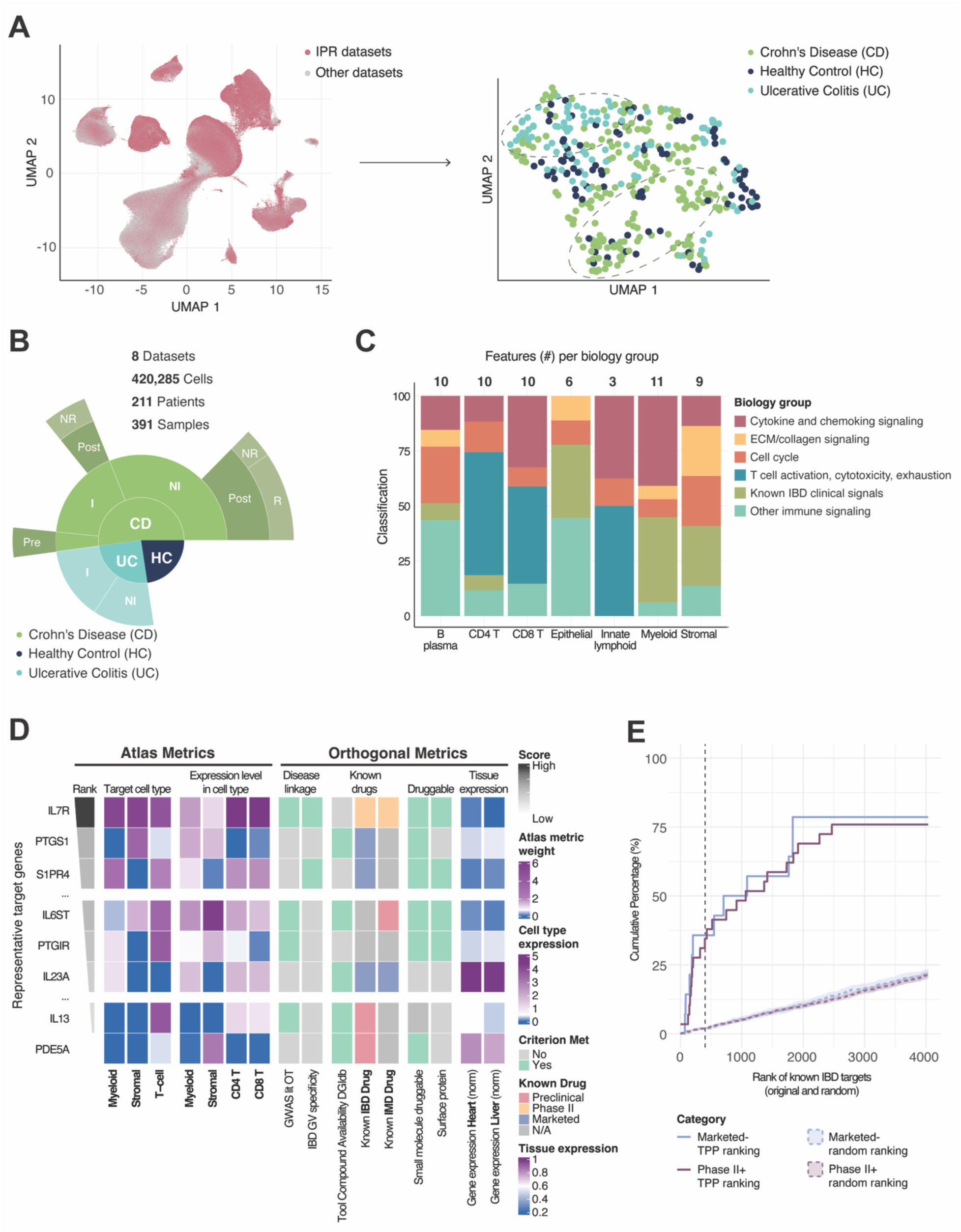
**Immune Patient Representation identifies disease-associated transcriptional programs and prioritizes atlas-derived targets.** A) The Immune Patient Representation (IPR) framework integrates single-cell data to identify cell-type-specific transcriptional programs distinguishing biological conditions. Left: UMAP of all cells in the IBD atlas, highlighting the cells from the 8 datasets used for IPR patient representation. Right: representative IPR embeddings showing sample-level structure across disease conditions. B) Composition of the 8 studies used for IPR analysis. Studies were selected to ensure comprehensive coverage of intestinal immune and stromal populations, sufficient patient representation across key disease contrasts, and to exclude sorted populations that could bias cell-type representation. C) Biological pathways enriched among significant disease-associated IPR features. The top identified IPR signatures were enriched in pathways related to known IBD clinical signals and cytotoxicity/exhaustion. D) Representative heatmap summarizing multi-criteria evaluation of selected atlas-derived targets. Columns display both quantitative and categorical prioritization metrics, including cell-type association, disease linkage, druggability, and tool-compound availability. Expression levels are shown as mean log-normalized values per cell type and across bulk tissue datasets (heart, liver) from AMICA DB^TM^. Details on the prioritization metrics are reported in Table S4. E) Distribution of known therapeutic targets across the ranked candidate-gene list. Targets are grouped by the highest clinical phase reached by the targeting therapeutic agent. Analysis included 14 genes with marketed targeting drugs, 2 genes with PIII targeted drugs, 36 genes with PII targeting drugs. CD, Crohn’s disease; HC, healthy control; UC, ulcerative colitis; I: inflamed, NI: non-inflamed, IMD: immune mediated disease.

To identify novel targetable disease mechanisms, biological questions were formulated as contrasts, including inflamed vs non-inflamed tissue, treatment-naïve vs post–anti-TNF samples, and responder vs non-responder groups (**Table S2**). IPR analysis produced a ranked set of 85 significant disease-associated features that was both cell type-specific and biologically interpretable **(Fig 2C and Fig S3B-H,** details in supplementary methods). These programs were distributed across a variety of cell types and pathway enrichment analysis revealed an overrepresentation of TNF signaling, cytotoxicity, and multiple known IBD-related clinical signatures (**Fig 2C**).

### Identification and prioritization of cell type specific therapeutic targets

We next used two complementary approaches to extract the most relevant candidate target genes contributing to each top feature. First, we ranked genes by the strength of their correlation with each IPR feature effect size value across patient samples using Pearson’s correlation. Second, we applied Median Absolute Deviation (MAD) normalization to the feature coefficients, to identify genes that exert unique, cell-type-specific influence within a given IPR. These two approaches served as a filtering step, yielding a pool of 4036 target genes, each associated with one or more cell type specific disease-relevant transcriptional programs within the IPR representation. Prioritizing from such a large gene list remains a critical challenge, as low success rates in drug development are frequently linked to the poor predictive validity of early-stage target selection^19,20^. To address this translational gap, we focused target prioritization on human-tissue derived, disease-relevant transcriptional metrics derived directly from the IPR model. This approach captures patient-derived, cell-type level variation that is typically overlooked by traditional approaches. These IPR based metrics included: 1) the strength of the IPR effect size, 2) the linkage of each IPR feature to inflammation and non-response contrasts, 3) the degree of differential expression across disease states, 4) involvement in network level pathways and 5) the association with cell types most strongly linked to disease-related transcriptional changes (**<u>Table</u> S4**).

To ensure alignment with our target product profile (TPP), favoring tractable, cell-type restricted, and potentially safer targets, we then integrated these IPR-derived metrics with publicly available criteria drawn from established principles in drug discovery^21–24^. These orthogonal metrics assessed novelty, genetic and functional linkage to disease, potential safety liabilities, and experimental feasibility (**Fig 2D, <u>Table</u> S4**).

We benchmarked this prioritization strategy by verifying the recovery of known targets of approved and late-stage clinical therapies for IBD. Among the IPR-derived genes approximately 50% of marketed drugs and 35% of targets from drugs in Phase II or III development were ranked in the top 400 genes (**Fig 2E**). These included genes such as TNF, integrin α4 (ITGA4), prostaglandin signaling (PTGS1/2), sphingosine-1-phosphate receptors (S1PR4), and JAK family kinases (JAK3), five of the six major mechanistic classes of current IBD therapies^2^. The sixth, IL-23, showed a robust IPR signal in myeloid and T cell subsets, but was deprioritized during TPP-based ranking because of its secreted nature and extra-intestinal expression (**Fig 2E**) illustrating the goal of our discovery framework: the identification of cell-type-specific, actionable, and translatable targets rather than broad, pleiotropic mediators. Overrepresentation analysis of known targets in the top 400 genes revealed a significantly higher probability of detection compared to random (Fisher exact test p<10^-6^). Overall, these findings confirm that the IPR-based discovery framework captures established therapeutic biology while enriching for novel, cell type-specific targets (**Fig 2D, E, Fig S4A**). Supporting this observation, approximately 15% of the top 400 genes overlapped with targets of drugs in PII or higher development for other immune-mediated diseases (**Fig S4B,C Table S5**), reflecting shared therapeutic mechanisms across multiple diseases.

Several targets in advanced clinical development ranked highly within our framework. IL7R emerged as the highest ranked gene, involved in disease associated pathways in multiple cell types including macrophages, fibroblasts, and T cells (**Fig 2D**). The clinical relevance of this finding is supported by recent promising Phase II results for the IL7R blocking antibody lusvertikimab in ulcerative colitis^25^, demonstrating the framework’s ability to contextualize known targets within cell-type-specific disease networks. The lower ranked genes included targets of drugs whose clinical development for IBD has since been discontinued, such as IL13 (tralokinumab^26^) and PDE5A (CEL-031^27,28^). While multiple factors can lead to drug discontinuation, this observation suggests that our transcriptional approach can effectively prioritize translatable targets and de-risk the drug discovery pipeline.

Having validated our framework, we focused on the top 400 ranked genes and applied a series of stringent filters to enrich for novel, actionable candidates for functional validation. Non-actionable genes included housekeeping genes, genes not amenable to *in vitro* validation, lowly expressed genes, and targets already in development for IBD. This process yielded 343 novel high-potential target candidates. The resulting target list is biologically diverse and includes genes involved in known IBD clinical pathways, cytokine and chemokine signaling, and pro-fibrotic signaling across multiple cell types (**Fig S3H**). Among the actionable targets, the largest number of candidates were associated with fibroblasts and mononuclear phagocyte subsets (**Fig 2C**, **Fig S4D**) providing a clear rationale for the development of functional validation models in these cell lineages.

### Identification and *in vitro* functional validation of PTGIR as a macrophage specific therapeutic target

Among the highly ranked genes, we identified PTGIR as a promising candidate for therapeutic blockade in IBD matching our desired target product profile (**Fig 2D**, **Fig 3A**). PTGIR lies within an IBD associated locus^5^ and encodes the G-protein coupled receptor Prostaglandin I2 Receptor. PTGIR acts as a receptor for prostaglandin I2 (PGI2) and, in the intestine, is expressed on the cell surface of myeloid cells, T cells, and stromal cells (**Fig S5A**). Despite the recent interest in the role of PTGIR in fibroblasts and T cell subsets, its function in intestinal macrophages, which are key regulators of inflammation and tissue remodeling, remains unknown^29,30^. Our IPR based analysis identified PTGIR as a top gene within transcriptional IPR features associated with several myeloid populations present in the inflamed intestinal tissue, including lipid-associated macrophages (LAM), and inflammatory macrophages (**Fig 3A**,**B****, Fig S5B**).

**Figure 3:**
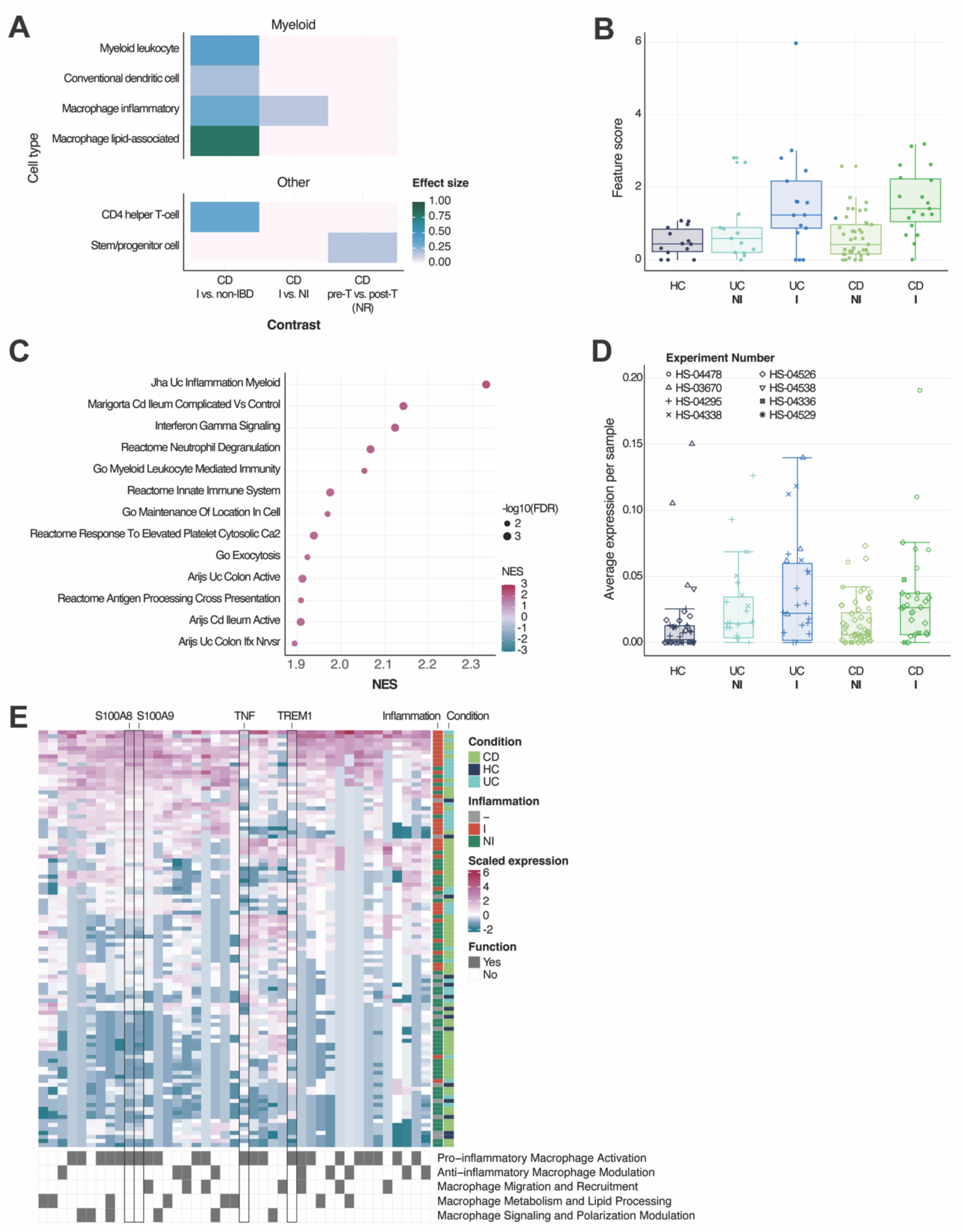
**A scRNA-Seq-based screening approach identifies PTGIR as a myeloid specific target in IBD.** A) PTGIR-associated signals from IPR-based clinically relevant contrasts. Rows represent cell types at the granular level; columns are the relevant disease and anti-TNF treatment-related contrasts. Color scale indicates the effect size of the IPR contrasts. B) IPR feature score for the top PTGIR-associated IPR feature (IPR feature 1) in total myeloid cells among inflamed and non-inflamed UC and CD samples, and in healthy controls. C) Gene set enrichment analysis (GSEA) for the LAM associated IPR feature 1. The normalized enrichment score (NES) for the top enriched features is shown. Dot size depicts -log 10 FDR (false discovery rate). D) Sample-level expression of PTGIR across disease groups in the IBD tissue atlas in total myeloid cells. PTGIR levels are shown as normalized sample pseudobulk expression. Data points are shaped by dataset of origin (Table S1). E) Heatmap of top AMICA-Reason^TM^ genes associated with the LAM associated IPR feature 1. Rows represent z-normalized pseudobulked expression of genes involved in one or more key biological processes, columns represent samples annotated by clinical metadata. Binary annotations to the right represent key biological process associations per gene. CD Inflamed n= 33, UC inflamed n=33, CD non-inflamed n=59, UC non-inflamed n=26, healthy controls n=43

Emerging evidence suggests that robust ML-approaches in single-cell analysis must pair computational inputs with biologically-grounded interpretability^31,32^. To investigate the translational relevance of the LAM-associated IPR feature, we employed two complementary approaches. First, we performed gene set enrichment analysis (GSEA) to test for the enrichment of a curated library of public pathways, as well as of gene signatures derived from orthogonal single-cell and bulk transcriptomics clinical IBD datasets. The enriched pathways reflected a pro-inflammatory transcriptional profile consistent with the innate immune activation observed in inflamed and treatment-refractory IBD^33–35^ (**Fig 3C, Table S1,S3**). However, enrichment analysis is not sufficient to extract coherent, context aware processes. Thus, we next used AMICA-Reason^TM^, Immunai’s agentic LLM-driven interpretation engine, to classify IPR gene-level signals into mechanistic modules in the context of IBD biology. AMICA-Reason^TM^ identified key inflammatory, chemokine-driven and metabolic modules up-regulated in inflamed vs non-inflamed and healthy intestinal samples (**Fig 3E**). The two subunits of the calprotectin heterodimer, S100A8 and S100A9, emerged as central drivers of this feature. PTGIR has been shown to control S100A8/9 expression^36^, linking the PTGIR-associated transcriptional network directly to a broadly used clinical biomarker of mucosal inflammation^37^. Together, the analysis of the LAM profile associated with PTGIR highlights metabolic and inflammatory programs consistent with the plasticity of macrophages in IBD. LAMs are increasingly recognized as a phenotypically plastic population whose function is modulated by environmental cues, but their contribution to IBD-pathology remains unclear^38^. Our findings suggest that activation of a PTGIR-associated network in LAMs contributes to disease-promoting pathways in IBD.

Consistent with a role in intestinal inflammation, the expression of PTGIR was elevated in inflamed CD and UC tissue compared to non-inflamed and healthy controls, a finding that was independently confirmed in a large independent single-cell cohort harmonized within AMICA DB^39^ (**Fig 3D, Fig S5F,G**). Analysis of non-intestinal expression within AMICA DB^TM^datasets (**Table S1**) confirmed the low expression of PTGIR in myeloid cells from the heart (**Fig S5C),** liver (**Fig S5D**) and systemic circulation (**Fig S5E)**, suggesting that myeloid specific targeting of PTGIR might have limited on-target off-tissue effects.

Given the strong association of PTGIR with a dysfunctional profile of inflammatory and lipid-associated macrophages, we next sought to validate its potential as a myeloid specific therapeutic target. We therefore developed an *in vitro* monocyte polarization model optimized to recapitulate the specific transcriptional states of these cells in inflamed IBD tissues. We screened over 30 distinct polarization conditions for human mononuclear phagocytes (MNPs) and used SystemMatch, a ML pipeline to select the model condition more closely resembling the target MNPs in our IBD single-cell atlas (**Fig 4A, methods)**^40^. The final conditions for functional validation were then selected based on the expression of PTGIR and its associated gene network ^29,36^. This analysis identified M-CSF+ IL1𝛽 as the ideal condition to test the role of PTGIR in lipid associated and inflammatory macrophages (**Fig 4B)**.

**Figure 4:**
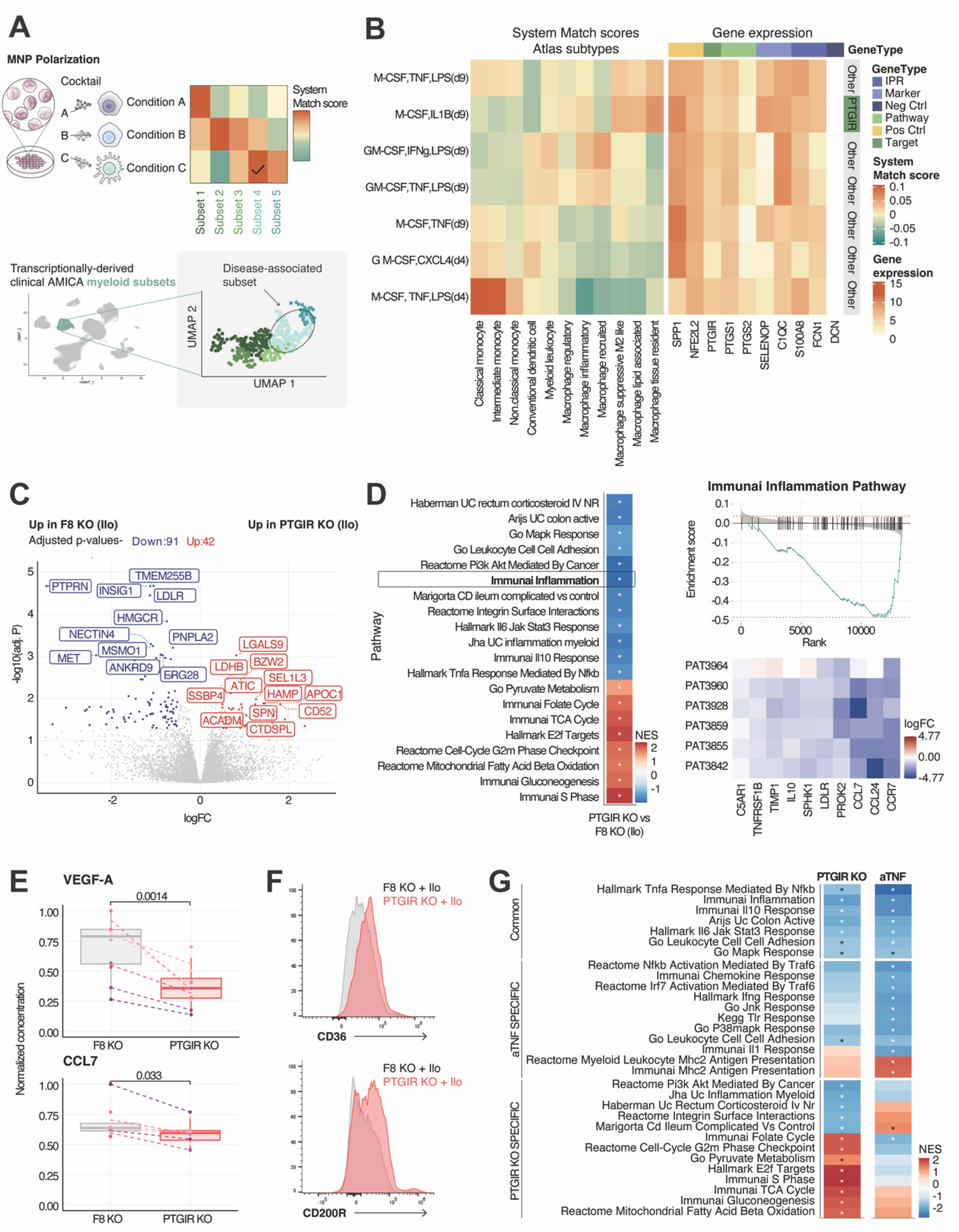
**Functional genomic validation of PTGIR in human mononuclear phagocytes** A) Schematic overview of the atlas informed model system selection matching disease-relevant cell populations to transcriptionally matched MNP polarization conditions. B) Atlas informed MNP polarization condition selection for PTGIR KO. The left heatmap shows the average z-score normalized SystemMatch values of transcriptional similarity to disease-relevant myeloid subsets including lipid-macrophages and inflammatory macrophages. The right heatmap shows the logCPM normalized expression of PTGIR, genes associated with lipid macrophages, and IPR feature 1 genes. Human monocytes across 6 donors were differentiated under multiple polarization conditions. Results for 6 representative conditions are shown. For the functional validation of PTGIR KO, KO of PTGIR and F8 (negative control) was performed in MNPs monocytes (n=5-6 donors). Cells were then cultured with M-CSF for 5 days, polarized with M-CSF and IL-1𝛽 for 2 days followed by stimulation with the PTGIR agonist Iloprost for 2 days (Ilo, 0.1nM, details in supplementary methods). C) Volcano Plot of differentially expressed genes in the PTGIR KO vs F8 KO contrast under iloprost stimulation. Top DEGs with |logFC|> 0.5 and adj_p < 0.1 are displayed. D) Left: Gene set enrichment analysis for PTGIR KO vs F8 KO contrast genes under iloprost stimulation. Right: Enrichment plot for Inflammation related genes and steroid refractory UC signatures^42^. The logFC expression of the top leading-edge genes is shown for each replicate in the 6 individual donors. E) LogFC for secreted VEGF-A (top) and CCL7 (bottom) upon PTGIR KO when compared to F8 KO following stimulation with Iloprost. Representative boxplot from 2 replicate experiments with n=6 each are shown. F) Representative flow cytometry plot of the surface protein expression of CD36 and CD200R in PTGIR KO and F8 KO controls under Iloprost stimulation. G) For the ex vivo comparison of the MoA of PTGIR and anti-TNF targeting, MNPs monocytes (n=5-6) were cultured with G-MCSF + IFN𝛾 + LPS for 6 days followed by treatment with a blocking anti-TNF antibody for 3 days (Infliximab, 1.25ug/mL before processing for multi-omic analysis (details in supplementary methods). Gene set enrichment analysis for the top pathways modulated by TNF blockade (anti-TNF vs unstimulated contrast) and/or PTGIR KO (PTGIR KO vs F8 KO contrast with Iloprost). Signatures are grouped as anti-TNF specific, PTGIR KO specific, or shared.

In this optimized MNP model, we used an endonuclease system to delete PTGIR and the negative control gene F8 across 6 healthy donors, achieving a median PTGIR KO efficacy of 68.32% with minimal donor-specific batch effects (**Fig S6A**). Bulk RNA-seq was performed and differential expression analyses used a donor-level paired design to separate KO and stimulation effects from donor-specific baselines (**Fig S6B**). Differential expression analysis revealed that stimulation with the synthetic PTGIR agonist Iloprost in F8 KO induced a strong pro-inflammatory transcriptional response (**Fig S6C-D**), reflecting changes observed in inflamed IBD tissues (**Fig S6D**). While PTGIR KO in unstimulated MNPs had a minimal transcriptional impact (**Fig S6E**), it reversed the pro-inflammatory state induced by Iloprost stimulation, causing a marked metabolic and transcriptional shift toward an anti-inflammatory phenotype (**Fig 4 C-D**).

Pathway analysis confirmed the suppression of inflammation associated pathways induced by PTGIR agonism, including disease associated signatures enriched in tissues from inflamed and treatment refractory patients (**Fig 4D**)^33,34,41,42^. PTGIR KO further led to the concomitant upregulation of pathways associated with oxidative metabolism and cellular stress recovery, such as TCA cycle and pyruvate pathways characteristic of suppressive macrophages (**Fig 4D)**. Consistent with this shift, analysis of the secreted proteome highlighted a decrease in pro-fibrotic and pro-inflammatory mediators such as VEGF-A and CCL7 (**Fig 4E)**. Flow cytometry further highlighted the increased surface expression of CD36 and CD200R previously associated with an anti-inflammatory/regulatory state (**Fig 4F)**.

We next compared the impact of PTGIR targeting to the mechanism of action of TNF blockade, a standard therapy for both UC and CD. As expected, *in vitro* TNF blockade in MNPs induced a strong anti-inflammatory shift with reduction of TNF response signaling and downregulation of disease-associated genes (**Fig, 4G**, **Fig S6F-I**). The pathways modulated by PTGIR KO included pathways distinct from those affected by direct TNF blockade *in vitro,* suggesting an orthogonal mechanism of action (**Fig 4G**). While both treatments downregulated the response to TNF and IL6 mediated signaling, PTGIR KO induced the downregulation of signatures associated with refractory and complicated IBD that were not affected by TNF blockade (**Fig 4D,G**). These findings position PTGIR as a promising macrophage-specific target capable of counteracting IBD associated pro-inflammatory pathways through mechanisms distinct from TNF antagonism.

### PTGIR deletion in macrophages reversed IBD disease signatures in the IBD atlas

To confirm the translational relevance of our findings, we next compared the transcriptional effects of PTGIR KO in macrophages with the cellular states observed in patient tissues. The transcriptional signature of PTGIR depletion highlighted a reversal of the pro-inflammatory IPR features associated with intestinal inflammation in myeloid cells, including those linked to lipid-associated and inflammatory macrophage subsets (**Fig 5A**).

**Figure 5:**
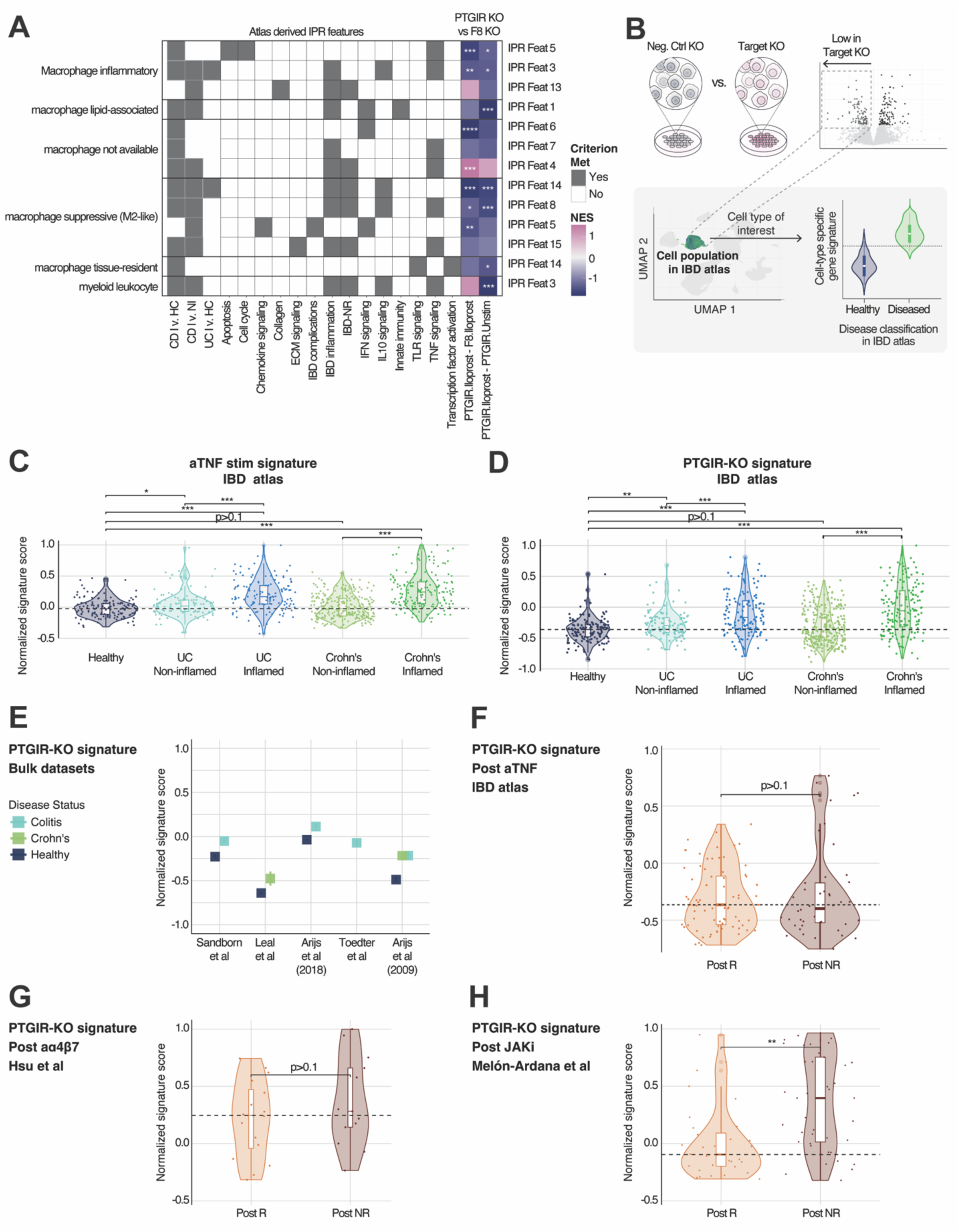
**clinical projection to clinical atlas confirms the relevance of PTGIR targeting and highlights differentiating mechanisms of action.** A) Heatmap of normalized enrichment scores for pro-inflammatory IPR feature gene sets in MNPs following either PTGIR KO or anti-TNF treatment in the *in vitro* MNPs model. B) Schematic overview of the clinical projection approach to validate the clinical relevance of the functional *in vitro* results. C) Projection of the gene signature downregulated in *in vitro* MNPs upon anti-TNF treatment to the IBD atlas, showing its enrichment in total myeloid cells stratified by disease group. D) Projection of the gene signature downregulated in *in vitro* MNPs upon PTGIR KO to the IBD atlas, showing its enrichment in total myeloid cells stratified by disease group. E) Projection of the gene signature downregulated in *in vitro* MNPs upon PTGIR KO to external bulk RNA-Seq datasets, stratified by disease group. F) Projection of the gene signature downregulated in *in vitro* MNPs upon PTGIR KO to the IBD atlas, showing its enrichment in total myeloid cells stratified by anti-TNF response groups. G) Projection of the gene signature downregulated in *in vitro* MNPs upon PTGIR KO to the Hsu et al dataset, showing its enrichment in total myeloid cells stratified by the anti-Integrin response groups. H) Projection of the gene signature downregulated in *in vitro* MNPs upon PTGIR KO to the Melón-Ardana et al dataset, showing its enrichment in total myeloid cells stratified by JAKi response groups.

We then applied a clinical projection approach (see Methods) that projects *in vitro* derived gene signatures on samples from the IBD atlas and independent validation cohorts, matched by cell type. This approach tests whether perturbing a target shifts disease-associated transcriptional programs towards those of the healthy or non-inflamed intestine (**Fig 5B**). As expected, *in vitro* TNF blockade with Infliximab (anti-TNF) treatment downregulated the disease-associated myeloid IPR features from the IBD atlas. These genes were consistently lower in healthy versus inflamed tissue across the atlas and in external validation cohorts (**Fig 5C, Fig S7A,B**), confirming that the clinical projection approach reliably captures clinically relevant therapeutic effects.

Consistent with the hypothesized role of PTGIR, treatment with its agonist Iloprost induced a strong pro-inflammatory transcriptional response, recapitulating transcriptional changes observed in inflamed tissues in the IBD atlas and in an independent single-cell dataset (**Fig S7C,D**). Conversely, projection of the signature downregulated upon PTGIR KO on the IBD atlas demonstrated that this set of genes was more highly expressed in macrophages from inflamed tissues compared with healthy or non-inflamed intestinal tissues (**Fig 5D,E**). These results suggest that targeting PTGIR can restore intestinal macrophages towards a non-pathogenic state, dampening disease-associated programs identified in inflamed tissues, a finding confirmed across multiple independent single-cell and bulk datasets (**Fig S7E,F**). To contextualize these effects within existing treatment paradigms, we compared the transcriptional phenotype downregulated by PTGIR KO with that induced by TNF blockade (infliximab), JAK inhibition (JAKi, tofacitinib)^43^ and anti-α4β7 integrin blockade (vedolizumab)^44,45^. The PTGIR KO signature was not significantly modulated by either anti-TNF or anti-α4β7 treatment, providing additional evidence that PTGIR blockade acts via orthogonal mechanisms to the existing biologics (**Fig 5F,G, Fig S7G-H**). In contrast, PTGIR KO showed significant overlap with the transcriptional response to JAK inhibition, specifically in treatment responders but not in non-responders (**Fig 5H**). This suggests that targeted inhibition of PTGIR might recapitulate anti-inflammatory effects of JAKi on myeloid cells while potentially avoiding off-target side effects associated with broad JAK inhibition^43,46^. Together, these results establish PTGIR as a macrophage-specific therapeutic candidate capable of reversing inflammatory gene programs in IBD and acting through pathways complementary to existing biologics.

### *In vitro* functional validation of IL6ST targeting demonstrates framework versatility in fibroblasts

Our discovery framework identified top candidates in MNPs and stromal cells. To demonstrate the applicability of our approach to non-immune cell types, we extended our functional validation to targets identified in fibroblasts that could play a key role in tissue remodeling and fibrosis. IL6ST (gp130), a signal transducer for the IL6 family^47^, was a highly ranked hit in stromal cells and T cells but not in myeloid populations (**Fig 2D, Fig S8A**). Its prioritization was further supported by GWAS evidence linking it to an IBD risk locus and a favorable safety profile due to low expression in cardiac and hepatic tissues (**Fig 2D, Fig S8B,C**).

Endonuclease mediated deletion of IL6ST in primary human fibroblasts, in the presence of one of its ligands, oncostatin M (OSM), caused a robust anti-fibrotic shift (**Fig 6A-C, Fig S8D**). This was characterized by the downregulation of pathways central to fibroblast activation, including TNF and TGF𝛽 responses and hypoxia signalling (**Fig 6A-C**) and decreased production of extracellular matrix proteins such as VEGF-A and Procollagen C-endopeptidase enhancer (PCOLCE) (**Fig 6D,E)**. Disease-associated signatures linked to complicated CD^34^ and non-response to anti-TNF and anti-integrin therapy were also reduced^35,48^ (**Fig 6C**). Together, these changes align with a shift away from pro-fibrotic, inflammatory fibroblast phenotype (**Fig. 6A-E**). In contrast, and consistent with atlas results, deletion of IL6ST in macrophages did not lead to a beneficial phenotype and rather triggered the upregulation of IFNγ-related pathways and robust secretion of inflammatory mediators (CXCL3, TNF, CCL15). (**Fig 6F-H, Fig S8E**).

**Figure 6:**
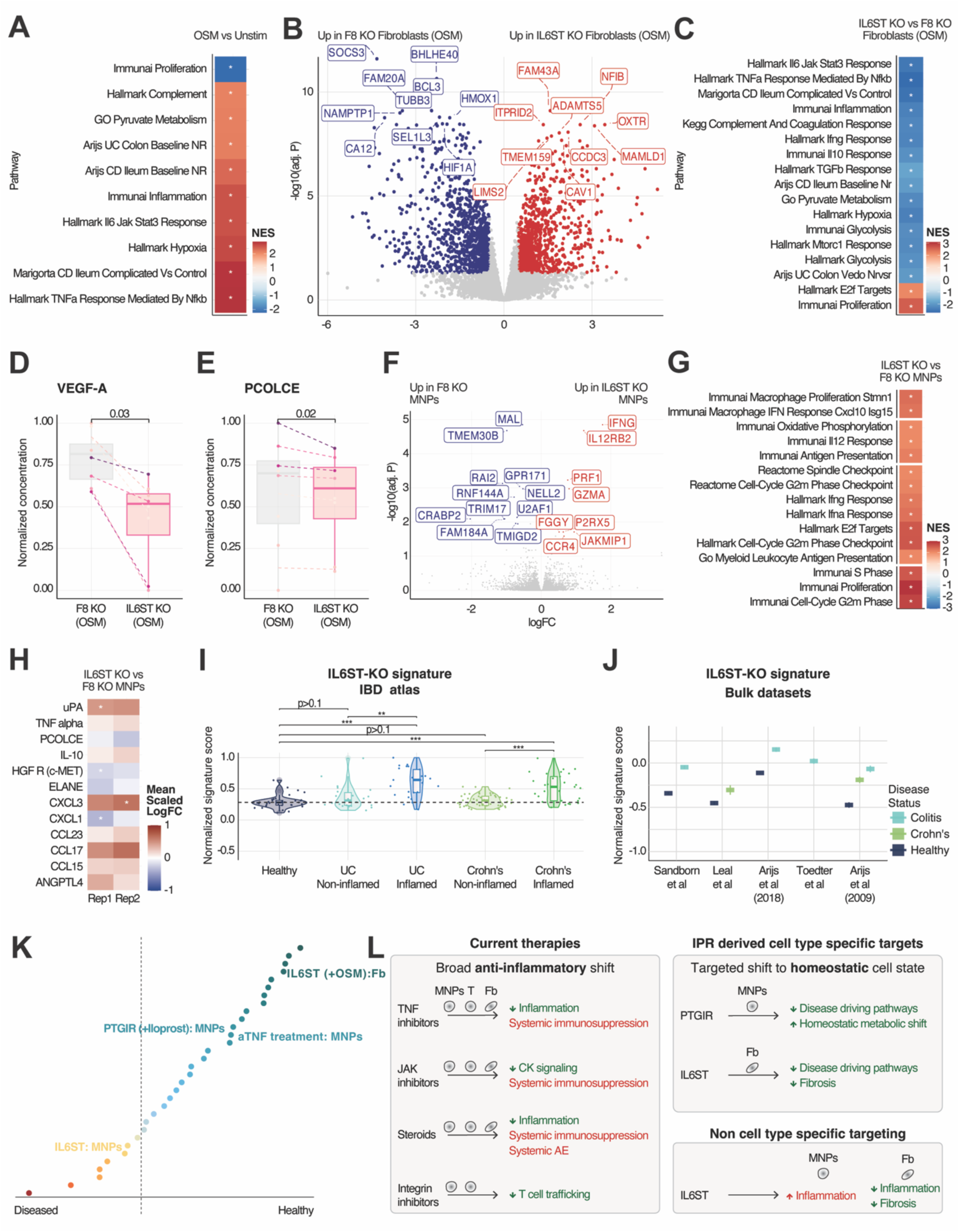
**IL6ST has opposing anti-fibrotic and pro-inflammatory functions in human primary fibroblasts and macrophages** For the functional validation of IL6ST in fibroblasts, KO of IL6ST and F8 (negative control) was performed in human primary small intestinal fibroblasts (n=2 donors) and, human primary colonic fibroblasts (n=2 donors). 6 days after KO, fibroblasts were stimulated with OSM (100ng/mL) and cultured for 3 days prior to multi-omic analysis (details in supplementary methods). B) GSEA analysis for the differentially expressed genes in the OSM vs untreated contrasts in fibroblasts. C) Volcano Plot of differentially expressed genes (DEGs) in the IL6ST KO vs F8 KO contrast in fibroblast in presence of OSM. Top DEGs with |logFC|> 0.5 and adj_p < 0.1 are highlighted. D) GSEA analysis for the DEG in the IL6ST KO vs F8 KO contrast in fibroblast in presence of OSM. E) Boxplot of the normalized supernatant concentration of VEGF-A in the supernatant of IL6ST KO and F8 KO in fibroblasts stimulated with the IL6ST ligand OSM. F) Boxplot of the normalized supernatant concentration of PCOLCE in the supernatant of IL6ST KO and F8 KO in fibroblasts stimulated with the IL6ST ligand OSM. For the functional validation of IL6ST in MNPs, KO of IL6ST and F8 (negative control) was performed in human monocytes (n=6 donors). Cells were then cultured with MCSF for 5 days, followed by polarization with TNF and LPS for 2 days. Cells were then stimulated with IL-6 (20ng/mL) for 2 days prior to collection. (details in supplementary methods). G) Volcano Plot of differentially expressed genes in the IL6ST KO vs Control F8 KO contrast in mononuclear phagocytes (MNPs). Top DEGs with |logFC|> 0.5 and adj_p < 0.1 are displayed. H) Heatmap of top GSEA of pathways modulated by IL6ST KO vs Control F8 KO in mononuclear phagocytes (MNPs) I) Top hits for multiplexed analysis of secreted proteins in the supernatant of IL6ST KO and F8 KO. J) Projection of the gene signature downregulated in primary human fibroblasts upon IL6ST KO to the IBD atlas, showing its enrichment in fibroblasts stratified by disease group. K) Projection of the gene signature downregulated in primary human fibroblasts upon IL6ST KO to external bulk RNA-Seq datasets, stratified by disease group. L) Comparison of IBD-atlas clinical projection effect scores for the differentially expressed gene sets generated by knockout of 19 candidate targets in MNPs and/or fibroblasts (Fb) with or without stimulation, with infliximab treatment in MNPs included as a positive control. Dashed line represents the inflection point where the effect scores become negative, indicating a more disease-like phenotype M) Summary scheme of the distinct mechanisms of action for PTGIR and IL6ST targeting based on their functional effect on MNPs and fibroblasts. Asterisks (*) denote significantly enriched pathways and significantly modulated analytes.

Clinical projection confirmed the clinical relevance of the functional validation results. In fibroblasts, the gene signatures downregulated upon IL6ST KO were consistently increased in inflamed CD and UC samples in both the IBD atlas and in external single-cell and bulk transcriptomic datasets (**Fig 6I, Fig S8F**) and bulk transcriptomic level (**Fig 6J**). Conversely, the pro-inflammatory transcriptional changes induced by IL6ST KO in MNPs clearly mapped to diseased tissue samples in the IBD atlas (**Fig 6K**).

Collectively, the clinical projection analysis highlights the cell type-specific therapeutic potential of PTGIR and IL6ST in macrophages and fibroblasts respectively (**Fig 6K**). KO of PTGIR in macrophages reflected a shift towards a healthy myeloid phenotype, matching the directionality seen *in vitro* with infliximab (anti-TNF) treatment (**Fig 6K)**. In contrast, IL6ST KO caused an anti-fibrotic shift in fibroblasts while its KO in macrophages triggered a disease-associated profile, suggesting that systemic or non-targeted inhibition of IL6ST would likely exacerbate inflammation (**Fig 6K)**.

Taken together, the functional validation of PTGIR and IL6ST highlights the strength of our atlas-based discovery framework to identify targets that contribute to disease-relevant pathogenic processes, together with the right cell population where this therapeutic modulation is beneficial, guiding precision strategies for patients with limited treatment options. (**Fig 6L)**.

## Conclusions

Novel target discovery in immune-mediated diseases remains challenging, as preclinical findings and functional readouts often fail to translate into clinical efficacy. This translational gap arises from the limited predictive validity of decision tools, insufficient human context during discovery, and suboptimal model selection^19,21^. Here, we establish a target discovery framework that addresses this gap by anchoring *in silico* discovery and *in vitro* functional validation to high-resolution, patient-derived single-cell data.

By leveraging the AMICA^TM^platform, and using a data foundation that harmonizes over 500 clinical samples in a single-cell IBD atlas of the intestine, we overcome the noise and heterogeneity of individual studies and achieve a high level of granularity in annotating the intestinal cell populations^13^. AMICA DB^TM^’s scalability enabled the integration of 4 additional datasets with over 150 samples that were used as external validation of the atlas derived findings. This integrated single-cell map of the healthy and diseased intestine provides direct human context for target discovery and model selection^9,10,13^. As the therapeutic landscape evolves, the AMICA^TM^ platform allows the seamless integration of new studies profiling the mechanisms of next-generation IBD therapies, maintaining an up-to-date clinical sample reference.

Our single-cell atlas of the human intestine forms the basis for our patient representation approach (IPR), a suite of built-for-purpose ML-tools that identifies disease-relevant transcriptional programs and the genes driving them within specific cell types. This single-cell anchored approach moves beyond the historical GWAS based identification of broad genetic loci, to pinpoint the specific genes, cell types, and cellular states driving pathology, providing a roadmap for the generation of precise therapeutic hypotheses for each novel target^21^.

This work demonstrates the power of our approach using public datasets and relatively simple ML methods. Of note, this scalable framework can easily integrate proprietary data in settings where high quality public data are sparse and incorporate more advanced representation-learning models to resolve rare, transitional or spatially restricted cell population signals, capabilities that are being pursued in our ongoing work.

Our model recapitulated established therapeutic pathways in IBD while uncovering new, cell-type aware therapeutic strategies. Our target prioritization strategy was designed to align with the desired target product profile, emphasizing tractable, cell-surface targets suitable for functional validation. However, these criteria are customizable, enabling application of the same framework to distinct therapeutic modalities or safety constraints. The strength of our target discovery framework is demonstrated by the *in vitro* validation of two cell type specific targets using atlas informed model systems. We show that targeting PTGIR leads to the metabolic reprogramming of macrophages and reduces their pro-inflammatory activity while targeting IL6ST reduces pro-fibrotic signatures in primary human fibroblasts, both via mechanisms orthogonal to those of existing biologics. The clinical projection of the *in vitro* signals to patient data, both within the IBD atlas and in external datasets, confirms that these targets can reverse disease-relevant cellular programs and suggests that their mechanisms are distinct from current therapies. Our target functional validation approach, which successfully recapitulates known biology of anti-TNF, can similarly be applied to define therapeutic mechanisms of action, stratify patients’ subgroups, and identify predictive biomarkers.

While our integrated approach establishes a powerful framework for novel therapeutic discovery, it is not without limitations. Our functional validation relied on primary human cell models that despite computational optimization, cannot fully recapitulate the complex environment of the diseased intestinal mucosa^49^. Nevertheless, clinical projection of the functional validation signatures confirmed the clinical relevance of our findings, suggesting that the main biological pathways are conserved in our models. As we look forward to further stages of drug development, our target discovery framework will extend to integrate more complex model systems.

Collectively, our work demonstrates how the integration of harmonized single-cell data, ML frameworks, and an agentic reasoning platform can overcome the long-standing challenge of extracting actionable biological insights from large-scale single-cell datasets^32,50^. Our next-generation framework is being expanded to integrate data from distinct modalities with the goal of achieving a cohesive understanding of disease biology and therapeutic mechanisms. These advanced models have the potential to facilitate novel precision medicine approaches by stratifying patient trajectories and identifying predictive biomarkers in a data-driven fashion. Collectively, these advances position the AMICA^TM^-IPR framework as a scalable and modular platform for data-driven drug discovery, transforming single-cell level insights into translatable therapeutic applications.

## Acknowledgments

The authors acknowledge Uzoma Nwankwo, for essential project management and operational support. We thank Hadass Jessel for expert assistance with figure design. We also acknowledge colleagues across Immunai and AstraZeneca for their valuable feedback and support.

## Credit Authorship Contributions

Anoushka Joglekar, PhD (Data curation: Equal; Formal analysis: Lead; Visualization: Lead; Investigation: Equal; Writing – review & editing: Equal), Ann Joseph, PhD (Investigation: Lead; Methodology: Equal, Formal analysis: Equal; Data curation: Equal; Investigation: Equal; Writing – review & editing: Supporting), Pavel Honsa, PhD (Data curation: Equal; Formal analysis: Equal), Klara Ruppova, PhD (Data curation: Equal; Formal analysis: Equal), Veronica Pizzarella (Investigation: Supporting), Amanda Honan, PhD (Investigation: Supporting), Devin Mediratta (Investigation: Supporting), Emily Vollmer (Investigation: Supporting), Evan Geller, PhD (Investigation: Supporting), Martin Valny, PhD (Data curation: Supporting; Formal analysis: Equal), Eva Macuchova, PhD (Data curation: Equal; Formal analysis: Supporting), Shiwei Zheng, PhD (Formal analysis: Equal), Alisa Greenberg, PhD (Formal analysis: Equal), Petr Taus, PhD (Formal analysis: Equal), Alina Kline-Schoder, PhD (Formal analysis: Equal), Renata Konickova, PhD (Data curation: Equal), Lucie Cerna, PhD (Data curation: Equal), Hila Sharim, PhD (Methodology: Supporting; Formal analysis: Supporting, Software: Equal), Lior Ness, PhD (Methodology: Supporting; Formal analysis: Supporting, Software: Equal), Giorgio Camilli (Investigation: Supporting), Eleni Chouri (Investigation: Supporting), Irem Kaymak (Investigation: Supporting), Joshua D’Rozario (Investigation: Supporting), Daniela Castiblanco (Investigation: Supporting), Joao Oliveira (Investigation: Supporting), Francesca Prandi (Investigation: Supporting), Nikolay Popov(Investigation: Supporting), Ana Laura Moldoveanu(Investigation: Supporting), Christopher Oliphant (Investigation: Supporting), Leire Escudero-Ibarz (Investigation: Supporting), Florian Uhlitz, PhD (Supervision: Supporting), Elizaveta Freinkman, PhD (Supervision: Equal; Project Administration: Equal), Jana Sponarova, PhD (Supervision: Supporting), Priya Vijay, PhD (Supervision: Supporting), Cailin Joyce, PhD (Supervision: Supporting), Irina Leonardi, PhD (Data curation: Supporting; Supervision: Supporting, Writing – Original Draft Preparation: Lead), Shikha Nayar, PhD (Supervision: Supporting; Formal Analysis: Equal, Project Administration: Equal), Adam Platt (Supervision: Supporting, Conceptualization: Supporting), Tatiana Ort (Supervision: Supporting, Conceptualization: Supporting), Greet De Baets (Supervision: Supporting, Conceptualization: Supporting, Project Administration: Supporting), Daniele Corridoni (Supervision: Supporting; Conceptualization: Supporting, Project Administration: Supporting), Aleksandra Wroblewska, PhD (Supervision: Equal, Writing – Review & Editing: Equal, Project Administration: Equal), Adeeb Rahman, PhD (Supervision: Lead, Conceptualization: Lead).

## Abbreviations

aTNF: anti–tumor necrosis factor antibody
CD: Crohn’s disease
DEG: differentially expressed gene
IBD: inflammatory bowel disease
IPR: Immune Patient Representation
LAM: lipid-associated macrophage
MNP: mononuclear phagocyte
scRNA-seq: single-cell RNA sequencing
UC: ulcerative colitis
UMAP: uniform manifold approximation and projection

## Disclosures

All authors declare potential competing interests. Authors affiliated with Immunai and AstraZeneca are employees of their respective organizations and may hold equity or stock options as part of standard compensation. The authors’ institutions have filed provisional patent applications related to aspects of the work described in this manuscript. No other financial, professional, or personal conflicts relevant to this manuscript are reported

## Data Transparency Statement

All clinical single-cell RNA sequencing and bulk transcriptomic datasets used for atlas construction and external validation are publicly available, with accession numbers provided in **Supplementary Table S1.** Data generated from *in vitro* functional validation experiments, including bulk RNA sequencing and associated analyses, are proprietary and are therefore not publicly available. Reasonable requests for methodological clarification may be directed to the corresponding author.

## Notes

### Summary of Updates

This version of the manuscript has been revised to update the authorship list.

