## Supplemental Files for "A ML-framework for the discovery of next-generation IBD targets using a harmonized single-cell atlas of patient tissue"

### Supplementary Figures

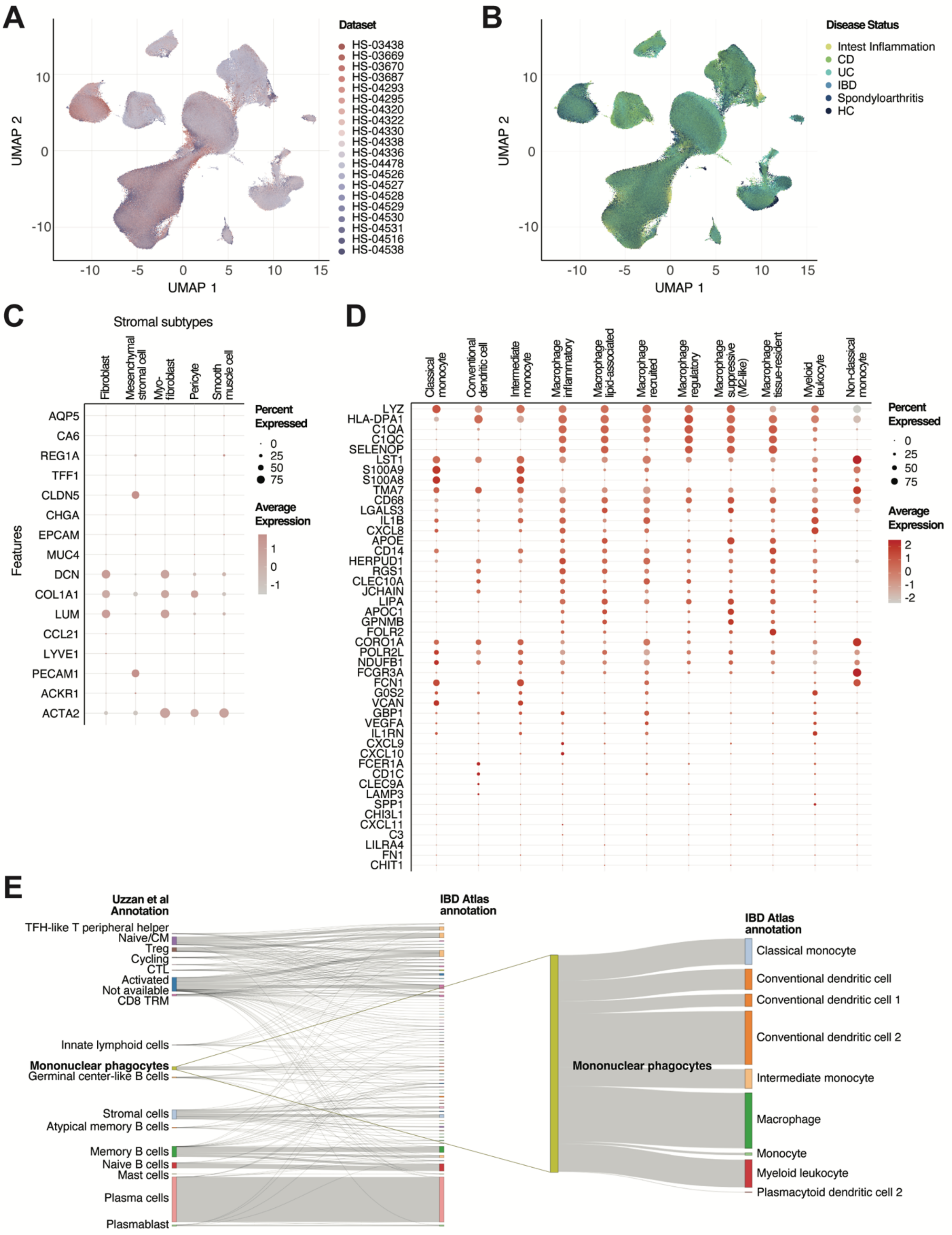


**Figure S1: Validation and characterization of the single-cell annotation**

A) UMAP visualization of the single-cell IBD atlas. Cells are colored by dataset. References and Source for each dataset in the IBD atlas are reported in table S1.
B) UMAP visualization of the single-cell IBD atlas. Cells are colored by disease status.
C) Dot plot showing the expression of canonical marker genes across annotated stromal cell subtypes in the IBD atlas. The size of the dot represents the percentage of cells expressing the gene, color indicates the normalized average expression level. The distinct expression patterns validate the accurate identification of stromal cell populations.
D) Dot plot showing the expression of canonical marker genes across annotated myeloid cell subtypes in the IBD atlas. The size of the dot represents the percentage of cells expressing the gene, color indicates the normalized average expression level. The distinct expression patterns validate the accurate identification of macrophage and monocyte populations.
E) Sankey plot comparing cell annotations from one of the original studies integrated in the IBD atlas (Uzzan et al., Nat Med, 2022)[1] with our reference-based approach, showing re-classification into higher-granularity subsets. The original annotation from Uzzan et al was replicated based on marker gene annotation of the clusters defined by the original manuscript. Activated: activated/effector T cell; Cycling: activated/cycling T cells; Naive/CM: naive/central memory; CTL: effector memory cytotoxic T lymphocytes. CD8 TRM: tissue resident memory CD8 T cells. Cells that could not be matched to the original annotation are labelled as not available. The IBD atlas reference-based approach annotates mononuclear phagocytes with higher granularity (right panel).
CD, Crohn's disease; HC, healthy control; IBD, inflammatory bowel disease; SpA, spondyloarthritis; UC, ulcerative colitis. UMAP, Uniform manifold approximation and projection

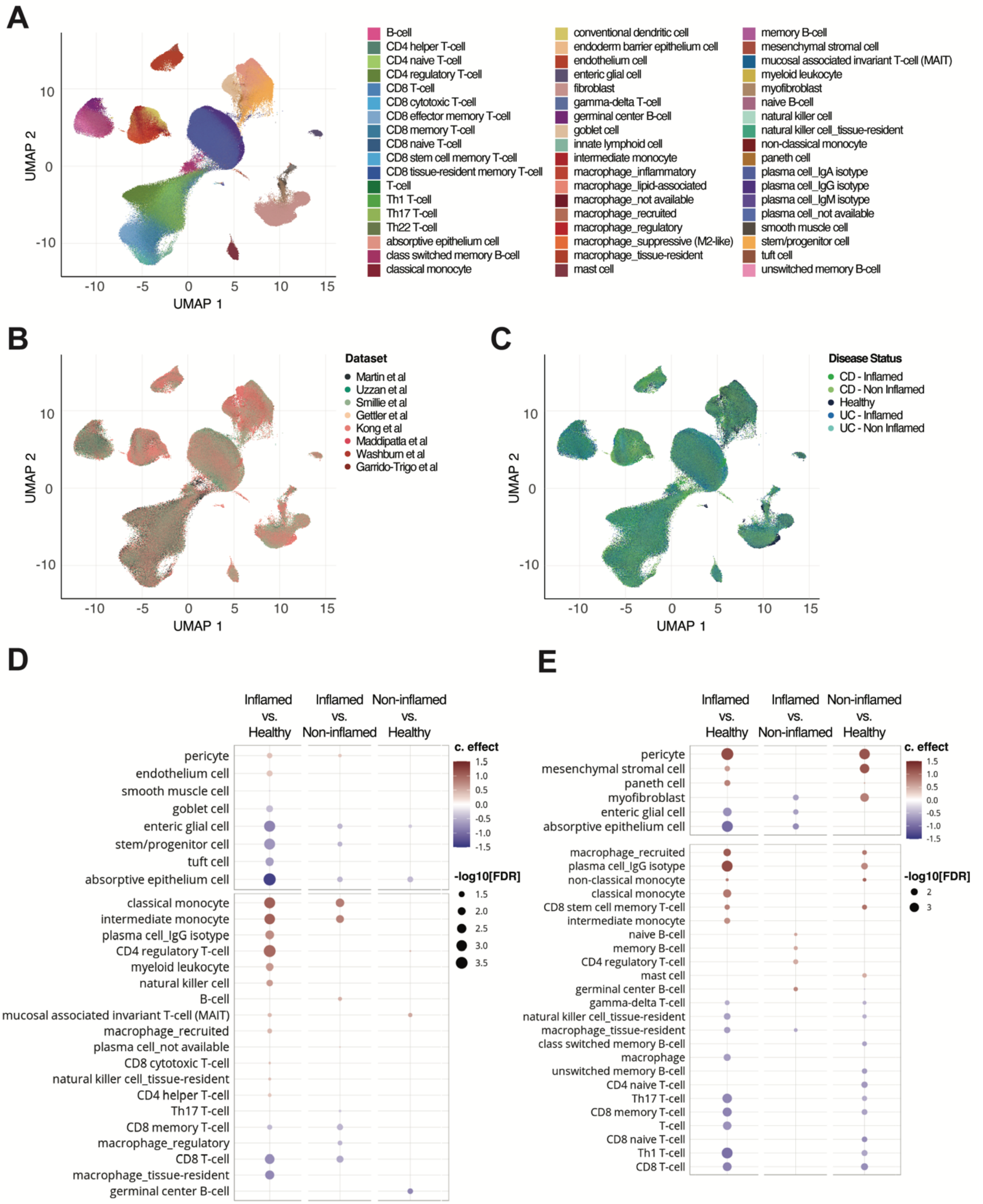


**Figure S2: Cell type level abundances per biological contrast in the 8 datasets selected for the IPR framework**

A) UMAP visualization of cells in the 8 single-cell datasets selected for sample representation, colored by cell type. B) UMAP visualization of cells in the 8 single-cell datasets selected for sample representation, colored by dataset; datasets are listed in Table S1.
C) UMAP visualization of cells in the 8 single-cell datasets selected for sample representation, colored by disease contrast.
D) Differential cell abundances across biological contrast in Crohn's disease. Size of dot indicates -log10 FDR while c.effect indicates the mean of the posterior distribution for each cell type abundance parameter.
E) Differential cell abundances across biological contrast in Ulcerative colitis. Size of dot indicates -log10 FDR while c.effect indicates the mean of the posterior distribution for each cell type abundance parameter.

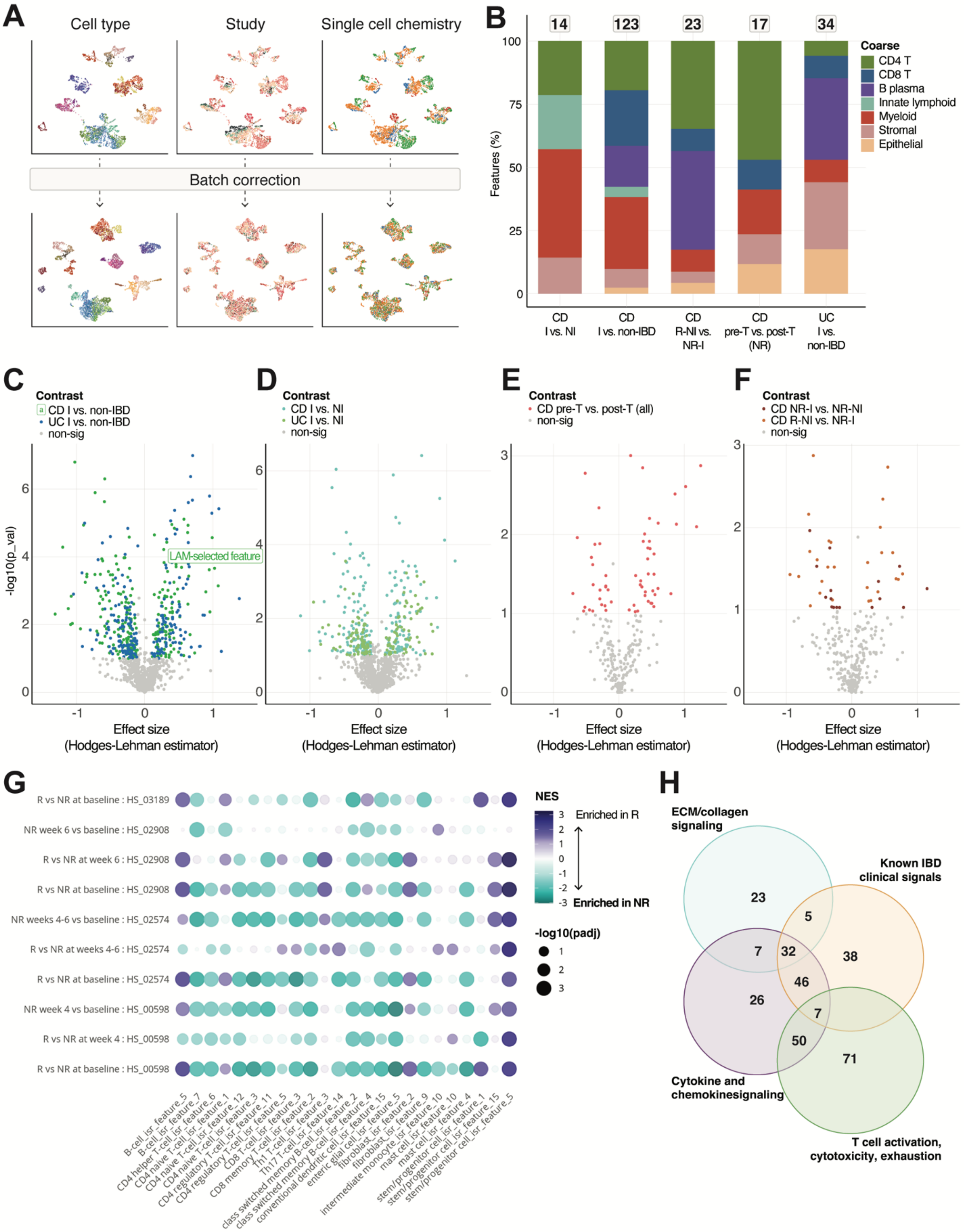


### Figure S3: Immunai Sample Representation

A) UMAP of the IPR atlas before and after batch correcting pseudobulked samples using a variational autoencoder. Each dot represents an individual sample. Samples are colored by cell type (left panels), study ID (middle panel), and single-cell chemistry (right panel).
B) Distribution of significant features across main cell type groups for each contrast category.
C-F) Hodges-Lehman estimator and p-values comparing latent space feature scores across all IPR features for the top contrast groups: disease (C), inflammation (D), treatment (E), Non response (F).
G) Enrichment of IPR features associated with non response in the IBD atlas across contrasts in bulk RNA-seq datasets (datasets are listed in Table S3).
H) Biological pathway associations for the top 400 candidate genes; genes are grouped by their primary IPR feature association. Known IBD clinical signals - clinically-derived signatures associated with anti-TNF NR or disease complications/ inflammation.

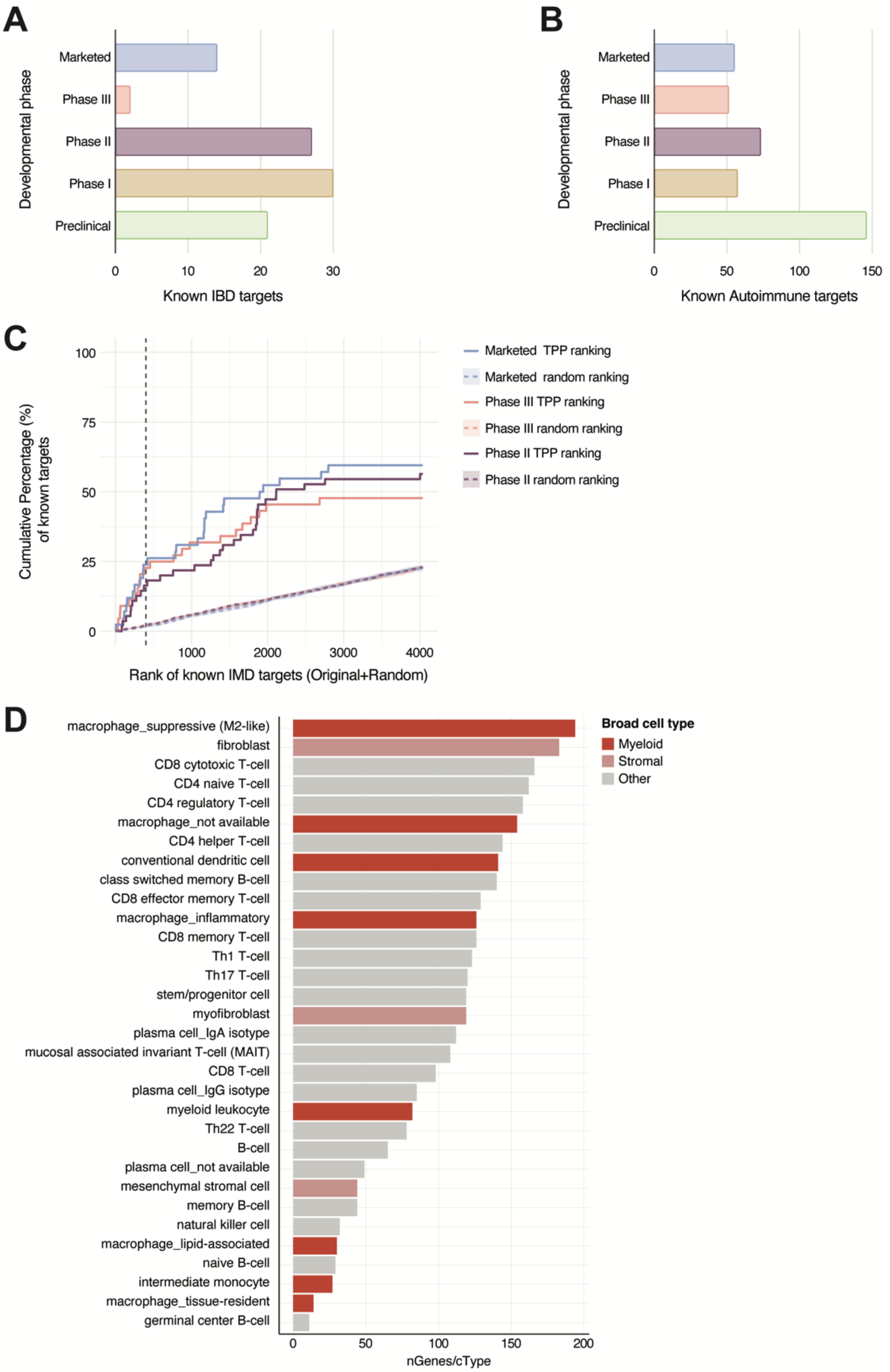


**Figure S4: Benchmarking and characterization of candidate therapeutic gene targets**

A) Developmental stages of gene targets for Inflammatory Bowel Disease (IBD). The figure shows the number of genes targeted by drugs for ulcerative colitis and/or Crohn's disease. For drugs no longer in development, the highest stage achieved is shown.
B) Number of genes with known targeting drugs in clinical and pre/clinical development for autoimmune diseases. Only genes targeted by drugs in active therapeutic development are shown. A complete list of autoimmune diseases considered this analysis is reported in **Table S5**.
C) Frequency distribution of successful phase I, II, III and approved targets for IMD among the ranked 4036 candidate target genes. Targets are grouped by the highest clinical phase reached by the targeting therapeutic agent.
D) Distribution of top 400 genes by cell type of origin. Each gene is counted multiple times if it was significant in multiple cell type-specific IPR features.

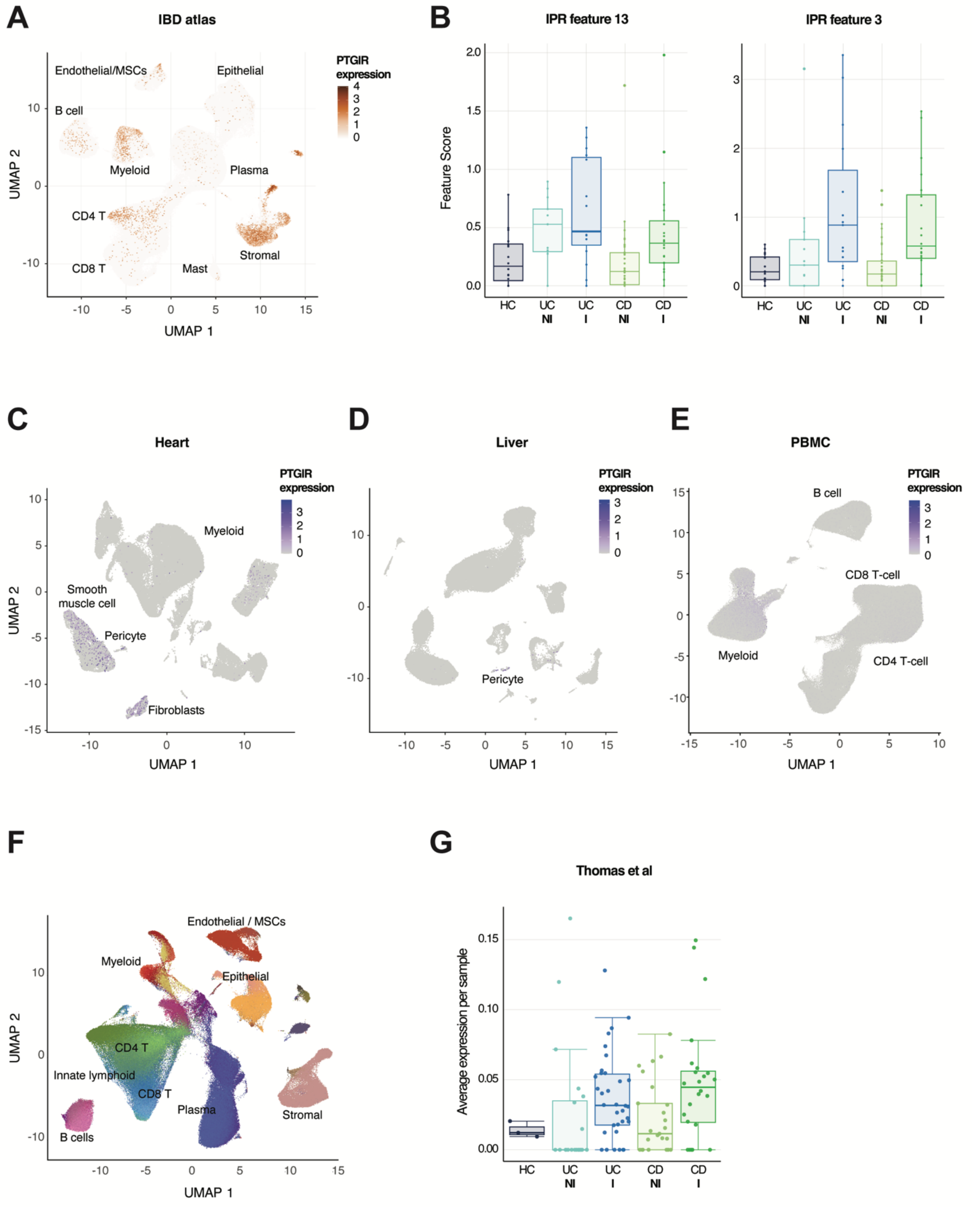


**Figure S5: Validation of PTGIR as a candidate myeloid target**

A) UMAP of cells the 8 datasets using for IPR sample representation, normalized expression of PTGIR is shown.

B) Expression of PTGIR associated inflammatory macrophages IPR feature 13 and myeloid leukocyte IPR feature 3 among total myeloid cells in the IPR atlas.
C-E). UMAP of PTGIR expression in healthy human heart (C),liver (D, and PBMC (E) single-cell datasets. Datasets are listed in Table S1.
F) UMAP visualization of the single-cells within the external validation Thomas et al dataset[2], colored by cell type.
G) Average expression of PTGIR in myeloid cells in the Thomas et al dataset[2].

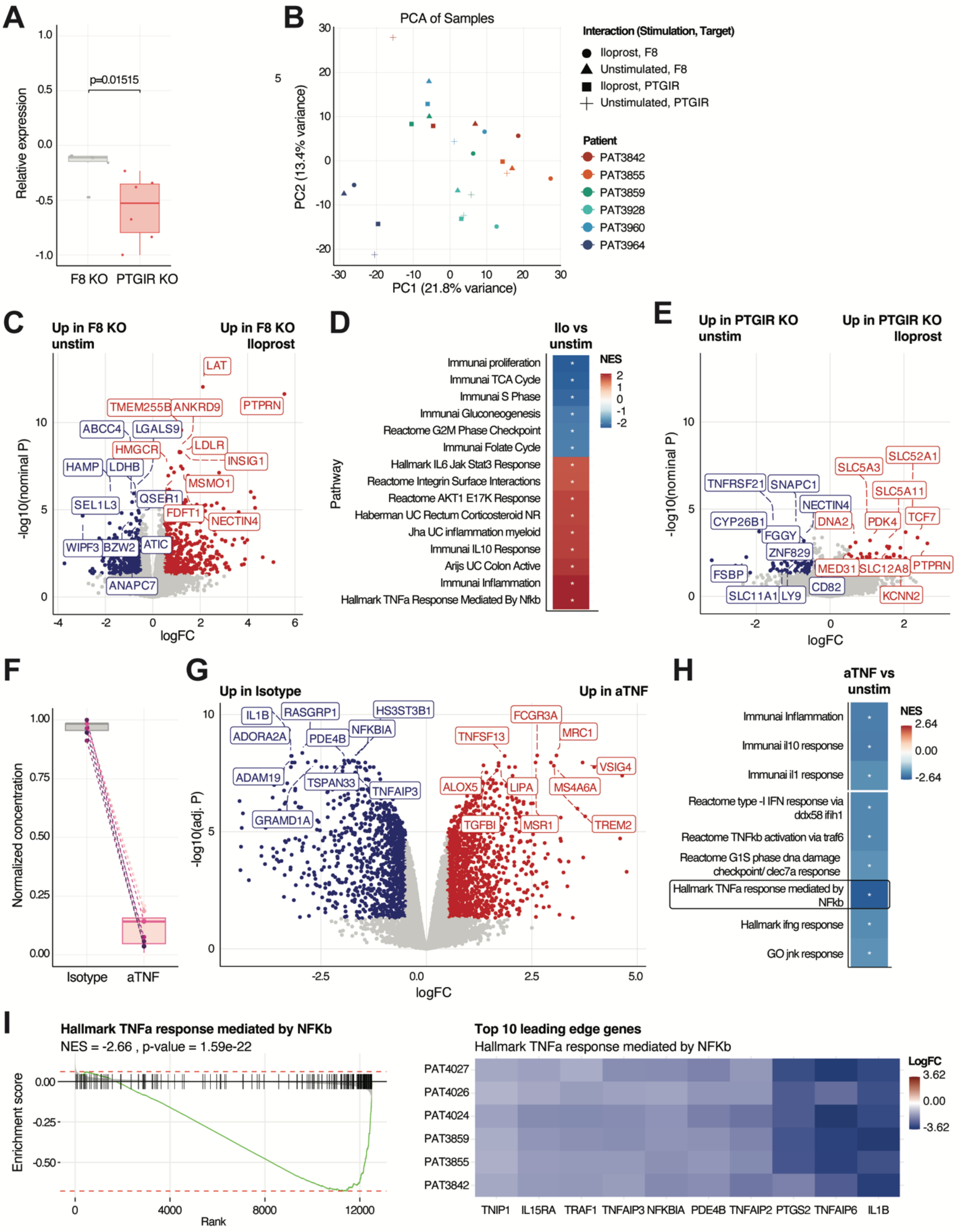


**Figure S6 Functional depletion of PTGIR in MNPs**

A) Validation of PTGIR KO efficiency in primary human MNPs. Representative qPCR quantification from 1 out of 6 PBMC donors showing relative PTGIR expression.

B) PCA of bulk RNA-seq data showing sample clustering across KO and stimulation condition. n=6 replicates per condition were used.
C) Differential expression for the iloprost vs unstimulated contrast in control F8 KO MNPs (n=6)
D) Gene set enrichment analysis for iloprost vs unstimulated contrast in F8 KO MNPs (n=6). Top enriched pathways are shown.
E) Differential expression for the Iloprost vs Unstimulated contrast in PTGIR KO MNPs n=6).
F) Quantification of soluble TNF in control F8 KO MNPs in presence or absence of the TNF blocking antibody infliximab.
G) Volcano plot for differentially expressed genes following treatment with Infliximab.
H) Gene set enrichment analysis for the infliximab vs untreated contrast in control F8 KO MNPs.
I) Gene set enrichment analysis for plot for Hallmark TNF response signature.

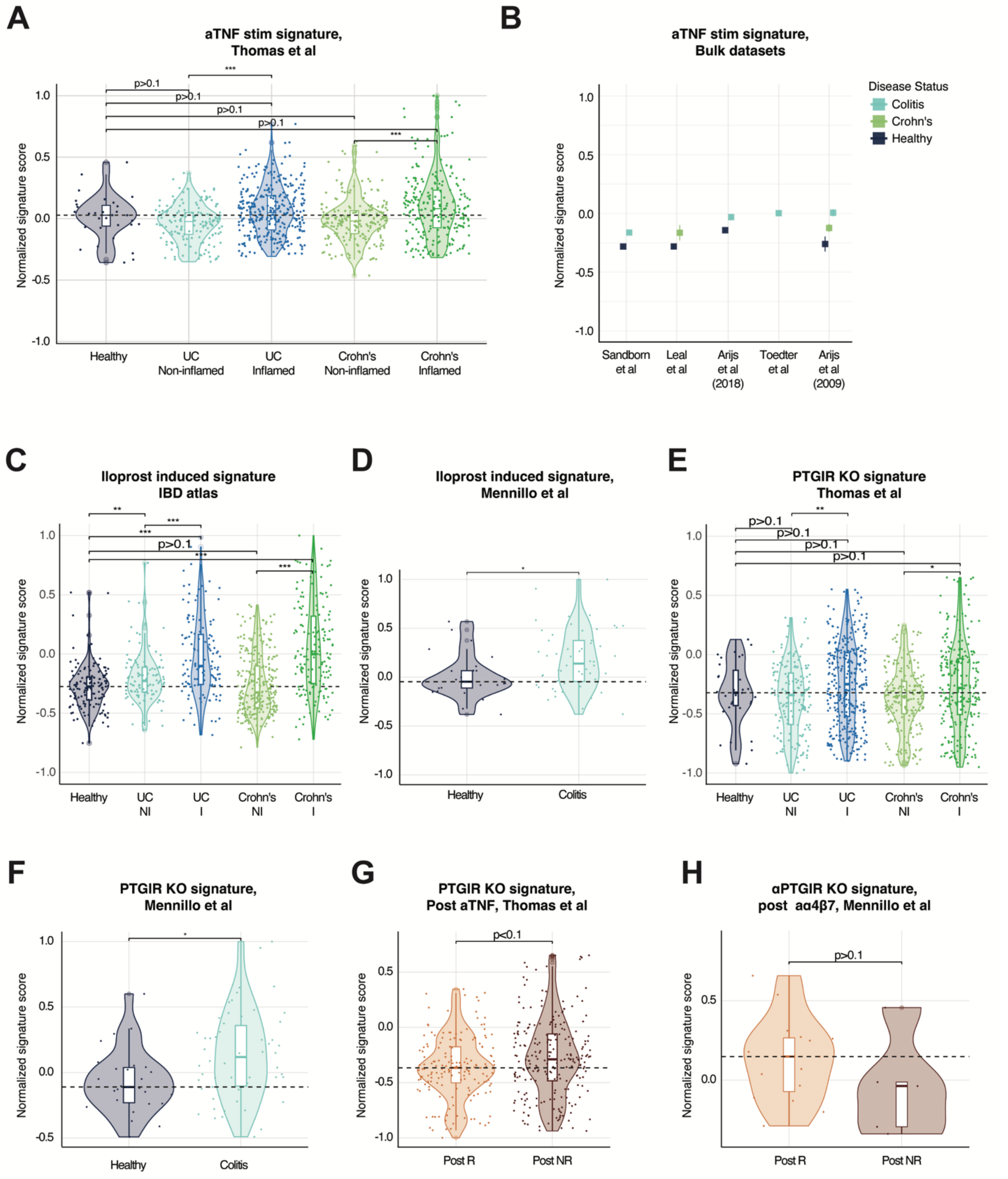


**Figure S7: clinical projection of PTGIR and TNF targeting signatures in human clinical samples**

A) Projection of the gene signature downregulated in *in vitro* MNPs upon anti-TNF treatment to the Thomas et al dataset, showing its enrichment in total myeloid cells stratified by disease group.
B) Projection of the gene signature downregulated in *in vitro* MNPs upon anti-TNF treatment to external bulk RNA seq datasets, stratified by disease group.
C) Projection of the gene signature upregulated in *in vitro* F8 KO MNPs upon stimulation with the PTGIR agonist iloprost to the IBD atlas, showing its enrichment in total myeloid cells stratified by disease group.
D) Projection of the gene signature upregulated in *in vitro* F8 KO MNPs upon stimulation with the PTGIR agonist iloprost to the Mennillo et al dataset[3], showing its enrichment in total myeloid cells stratified by disease group.
E) Projection of the gene signature downregulated in *in vitro* MNPs upon PTGIR KO to the Thomas et al dataset, showing its enrichment in total myeloid cells stratified by disease group.
F) Projection of the gene signature downregulated in *in vitro* MNPs upon PTGIR KO to the Mennillo et al dataset, showing its enrichment in total myeloid cells stratified by disease groups.
G) Projection of the gene signature downregulated in *in vitro* MNPs upon PTGIR KO to the Mennillo et al dataset, showing its enrichment in total myeloid cells stratified by anti-integrin response groups.

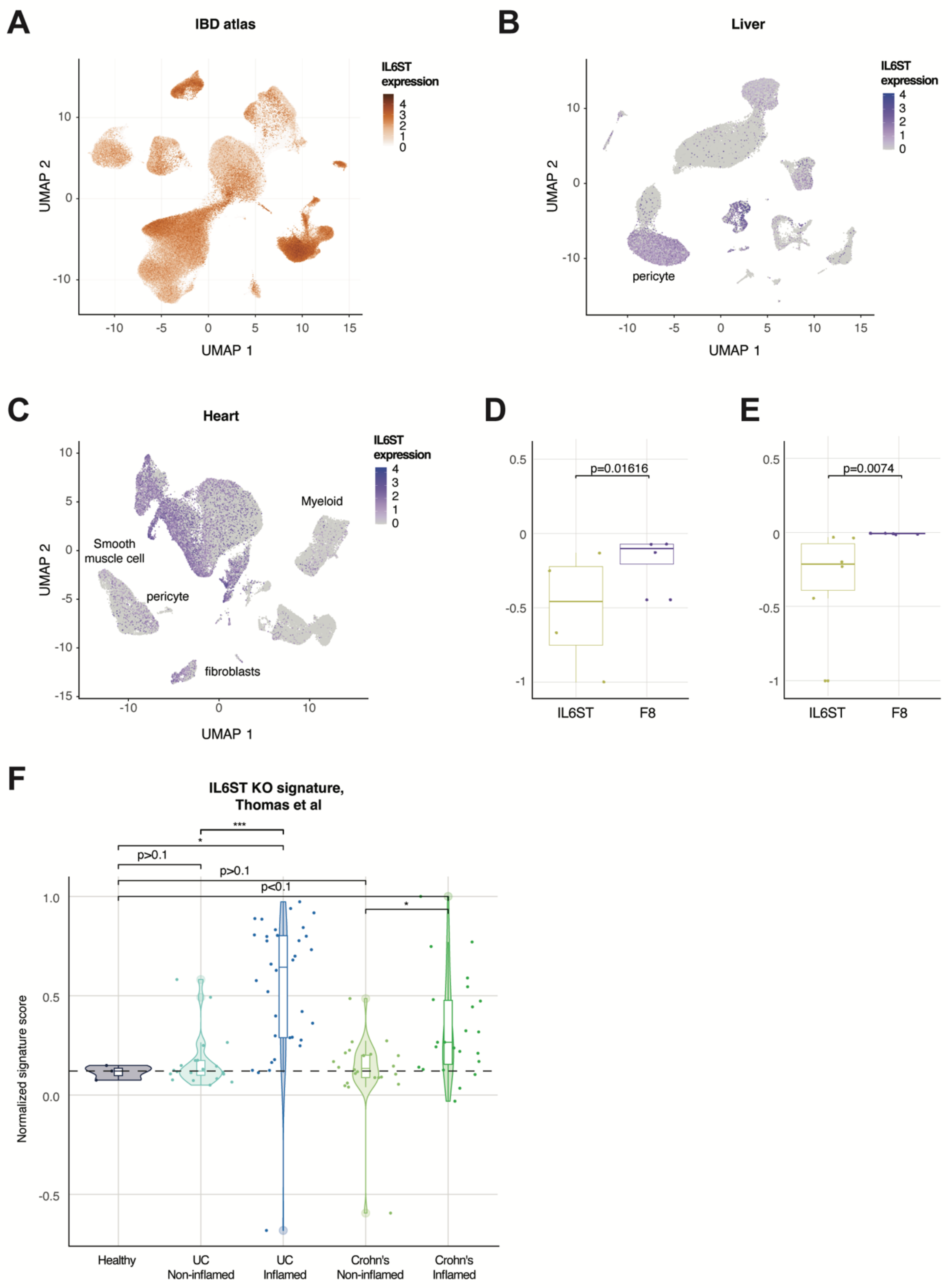


**Figure S8: Functional validation and clinical projection of IL6ST**

A) UMAP of IL6ST expression in the IBD BD atlas.

B) UMAP of IL6ST expression in the liver.

C) UMAP of IL6ST expression in the heart.
D) KO efficacy of IL6ST in primary human fibroblasts.
E) KO efficacy of IL6ST in primary human MNPs.
F) Clinical projection signature score for the *in vitro* downregulated genes following ILS6T KO in fibroblasts to the disease groups within the Thomas et al dataset[2].

### Supplementary Methods

#### IBD tissue atlas construction

All datasets used to construct the intestinal atlas were retrieved directly from AMICA DB™, Immunai's harmonized repository of proprietary and public single-cell studies. The AMICA™ plattform continuously ingests and processes scRNA-seq datasets through a standardized workflow, enabling consistent quality control, normalization, and annotation across studies (**Table S1**). Based on a review of existing literature, 20 single-cell studies relevant to IBD comprising 530 samples from 229 patients spanning multiple 10X chemistries were selected for atlas construction (Supplementary Table **1**). Samples not derived from primary gut tissue were excluded (e.g., PBMCs, organoids), sorted populations of non-immune cells (e.g., vascular cells, iPSC), and studies using non-UMI based library types (e.g., Smart-seq, SMARTer, Seq-well). To ensure the quality and robustness of the atlas, studies with fewer than 1000 cells or 2 samples, or those lacking available raw data, were also excluded. Raw data and clinical metadata were uniformly integrated in AMICA DB™ using our QC and harmonization pipeline. Raw data was processed internally using 10x Genomics Cell Ranger v5.0.1 to obtain count matrices. Filtering thresholds standard to single-cell data processing were applied to remove low-quality cells, wherein cells with at least 200 genes, 300 UMIs, and less than 10% of UMIs arising from mitochondrial genes were excluded. Additionally, cells above the 95th percentile of UMIs and 99th percentile of gene counts were excluded as potential homotypic doublets. The integrated data was processed using Seurat (v4.1.1), normalized using scTransform (v0.3.3), and corrected for sample batch effects using Harmony (v1.0). The resultant cell clusters were further clustered to obtain fine-grained cell subtypes and cell states, which were the annotated leveraging our proprietary cell-type ontology and tissue reference. Furthermore, the metadata from all 20 studies was downloaded, expertly curated by annotating with controlled vocabularies and annotation enrichment, extensively reviewed, and integrated into the final Seurat object containing 994,206 cells across 129 cell subtypes and cell states.

#### Differential cell type composition analysis

To identify cell types with significant changes in abundance between conditions, we performed a differential composition analysis using the sccomp package[4]. The "c effect", represents the mean of the posterior distribution for each cell type abundance parameter, and was used as a surrogate for the overall effect size on cell type proportions within each contrast[5].

#### IPR construction

Samples for this analysis were selected from eight studies to ensure a comprehensive representation of intestinal cell populations and sufficient sample numbers across key biological contrasts (Table S1).
Studies were excluded based on the following criteria: 1) Cells were sorted for specific subpopulations prior to scRNA-Seq analysis (n=10), as batch correcting sorted and unsorted samples could introduce technical artifacts. 2) The study involved a non-IBD disease state (immune checkpoint colitis; n=1). 3) Cells could not be assigned to requisite patient clinical metadata (n=2).
The resulting IPR was constructed from 8 high quality single-cell studies (Table S1). For the IPR model used here, for each cell type, we generated pseudobulk expression profiles, performed linear dimensionality reduction, and utilized the gene compositions of the dimensions as gene sets for further analysis. The dimensions of the latent space, non-inflamed, anti-TNF responder vs. non-responder) using a Wilcoxon

rank-sum test. Features with a p-value < 0.1 were considered significant. To ensure clinical relevance, gene signatures calculated from the gene components of IPR features distinguishing anti-TNF response were validated for enrichment in external bulk transcriptomics cohorts; non-validating features were excluded from further analysis (**Fig S3G**).

#### Identification of disease-driving transcriptional features

To identify disease-driving transcriptional features, we systematically evaluated their performance across 12 biological contrasts of interest (Table S2). For each contrast and cell type, a minimum cutoff of >= 3 samples per group was required, therefore some cell types were dropped for some contrasts. For each contrast, a Wilcoxon rank-sum test was applied to every feature, defined as the combination of IPR latent component and cell type. Features were then ranked by effect size (location difference, representing the median difference between the two groups) and p-value. A positive location_diff means the feature is higher in the second group in the contrast (e.g. CD I vs. non-IBD). A negative location_diff means the feature is higher in the first group in the contrast (e.g. CD I in the same example) Significance threshold for features was set at p < 0.1 and identified 92 key cell type-specific transcriptional features relevant to intestinal inflammation and anti-TNF non-response

Validation in bulk RNA sequencing datasets

24 IPR features separated anti-TNF NR from R with sig threshold of p < 0.1. Gene signatures from these features were tested against external bulk transcriptomics cohorts (Table S3). 18 of these 24 features (75%) were confirmed to be enriched in non-responder samples, validating the discoveries from our IPR atlas. The 7 IPR features with a discrepant directionality between the IPR atlas and the bulk RNA cohort were excluded from further analysis (**Fig S3G)**. Any gene that was in a feature that did not validate in bulk transcriptomics cohorts (N=314 genes from N=7 features) was filtered out and not incorporated into the final long list of candidate genes.

#### Interpretation with AMICA™-Reason

Immunai's agentic AI platform for structured biological reasoning, AMICA-Reason™ was used to systematically contextualize the biological function of the top cell type-specific IPR features associated with disease and inflammation. The AMICA-Reason™ framework is composed of several reasoning modules, operating at nested biological resolutions, to enable molecular, cellular, and systems-level interpretation. AMICA-Reason™ employs an orchestration framework that coordinates multiple LLM-powered reasoning agents (e.g. OpenAI or Google) and chain-of-thought prompting to generate, contextualize and evaluate mechanistic immune hypotheses. Overall, this agentic reasoning framework enables systematic, robust and scalable interpretation of complex biological data, turning model predictions into testable mechanistic hypotheses. AMICA-Reason™ enabled A) Functional characterization of gene sets, B) Cellular insight generation.

A) Functional characterization of gene sets. This module applies agentic reasoning to characterize the biological functions represented within a single Immunai Sample Representation, IPR, feature. This module uses two distinct components:
   a) LLM-based functional analysis agent: an LLM extracts the biological roles of each gene within a given gene set. Genes involved in related biological processes are grouped together and assigned a function. The analysis integrates public biomedical sources (e.g. Human Protein Atlas, PubMed) with the specific biological context of the contrast (e.g.,

blood, SCLC, monocytes). Through iterative reasoning and contextual reflection, the module refines its functional annotation to ensure coherence and biological plausibility.

b) Enrichment analysis: an over-representation statistical analysis of the gene list is then performed against Immunai's proprietary molecular signature database. The signature database is a compendium of proprietary signatures generated from Immunai experimental or curated public AMICA™ data. Statistically significant signatures enrich the LLM-derived functions. This step provides an orthogonal, data-driven validation that enhances confidence in the inferred biological functions.

B) Cellular insight generation. This module describes the immune processes within each cell type that are associated with the biological contrast of interest. The biological functions for features calculated by the first module are fed into an LLM-powered reasoning agent that synthesizes and evaluates shared biological themes. Specifically, the structured agentic reasoning combines generation, reflection, refinement and iteration to extract common immune processes across features. The LLM identifies related immune functions that recur across features and gene sets and evaluates each function's biological meaning (e.g., T cell activation vs. T cell suppression) as well as its direction in the data (e.g., up or down in IBD). Only features, functions, and processes that are biologically related and shifting in the same direction for a given contrast are grouped together. Through an automated iterative process, the agent can detect and resolve apparent contradictions, e.g., by digging deeper into supporting evidence and refining the functional annotations (e.g. cytotoxicity or effector functions annotated as both up- and down-regulated in a certain cell type, can be refined to "enhanced granule-mediated target cell killing", supported by upregulation of GZMA, GZMK, and "impaired adhesion-dependent co-stimulation" supported by downregulation of CD226, ITGA1, and NCAM1). True opposing functions that can arise from pleiotropic responses within a cell type (e.g., pro- and anti-inflammatory) and could not be resolved by the agent, are kept separate. The result is a collection of functionally refined, well-supported insights that characterize the most significant immune processes within each cell type, relevant to the biological question of interest.

#### Differential expression analysis

Differential gene expression (DEG) was performed by pseudo-bulk analysis. Gene counts were aggregated (sum) for each sample (per patient and time point) and cell type. If a sample per cell type had less than 15 cells, that sample was removed from the analysis for that cell type. If the analysis included e.g. testing for post-treatment effects while normalizing for baseline, patients for which we did not have the matched samples (pre- and post-treatment) were further excluded from the analysis. Differential expression was performed with the limma-voom R package (version 3.42.2). To normalize for inter-patient variability at baseline when performing DEG analysis for treatment effects, patient ID was added as a covariate to the model. 10 genes that were not captured in IPR transcriptional features were identified by this method and added to the candidate long list.

#### Extraction of candidate target genes from transcriptional features

Each top IPR feature (as determined by Wilcoxon rank sum test against a contrast) was distilled into a ranked list of candidate genes. Genes were included if they met one of these 2 criteria:

1. Pearson's correlation between gene expression and IPR feature value >=0.6
2. Pearson's correlation between gene expression and IPR feature value >=0.3 and Median absolute deviation (MAD) normalization of IPR features coefficients >= mean + 1.5sd

#### Orthogonal metrics for gene discovery prioritization

Candidate target genes were ranked by integrating the IBD atlas derived transcriptional evidence with a wide range of orthogonal, publicly available data (**Table S4**). Atlas-derived metrics (IPR feature strength to disease associations, DEGs, cell-cell communication patterns), and orthogonal metrics were aggregated towards ranking targets using a weighted approach. For each gene candidate, each metric value was either min-max normalized or encoded as binary and aggregated using a weighted sum. Weights were assigned by priority level with atlas-derived IPR and DEG metrics contributing most heavily, followed by genetic evidence, small molecule druggability, network-topology, known targets, heart/liver gene expression, and mouse evidence (**Table S4**). This resulted in a ranked list, from which 20 were validated through Endonuclease-mediated knockout.

Cell-cell communication

Significant cell-cell interactions were identified using the MultiNicheNet package: https://github.com/saeyslab/multinichenetr . This allows for prioritization of LR pairs based on their expression and prioritization based on ligand activity inferred from downstream signaling of LR interactions. The output is an aggregated prioritization score (0-1). We considered any LR prioritization scores of >0.6 as important to include in our ranking approach.

Protein-protein interactions

Protein-protein network data was obtained from the STRING database (Accessed Dec 2023) after removal of low confidence interactions (combined confidence score > 500)[6]. The combined confidence score was computed by combining the interaction probabilities from the different evidence channels and used as edge weights to give more attention to interactions with stronger evidence. Highly variable genes expressed in each major cell-type were used as edges. As expected, we verified that known IBD and IMD drug targets have higher node degree and betweenness centrality, and to have lower topological coefficient rendering their targeting less susceptible to network robustness and less prone to compensation through redundant signaling mechanisms[7].

IBD genetic evidence

Candidate targets are evaluated for presence of known genetic association by cross-referencing with a curated list of IBD GWAS signals and gene-disease relationships (supp table). We also utilized Genetic Target/Disease evidence from the Open Targets (OT) Database[8] (sum of scores > 0.5). In addition, we used Locus2Gene[9], a ML method to identify the most likely causal genes by integrating and summarising the effect of variants (GWAS Catalog and UK Biobank) based on genetic and functional genomic data. L2G score can be interpreted as a probability of a gene being causal. These were utilized to score each gene for IBD genetic evidence.

Known IBD and IMD drugs

Candidate drug targets were evaluated for targetability by known drugs for IBD and immune mediated diseases (IMD)-drugs by cross-referencing with a curated list of drug-targets. Drugs at both preclinical and clinical stages as well as marketed drugs were included. IMD diseases queried are listed in Table S5

Toxicity via heart and liver expression

1 heart and 6 liver scRNA-seq public studies were processed, curated, and harmonized within AMICA™ DB (Table S1). Gene expression was evaluated in all 7 studies, and each gene was scored for whether it appeared in the top 25% or bottom 25% of expression in these tissues.

Mouse immune and toxicity evidence

Immune: Candidate Immunai target genes are evaluated for association with immune/intestinal phenotype in mice via the IMPC mouse phenotype database[10] (Data release 20.1). Genes are encoded as "yes" if the following phenotypic abnormalities are reported as significant: Intestine: rectum, colon, cecum, ileum, duodenum, jejunum abnormalities. Immune: any immune cell abnormality

The IMPC mouse phenotype database was also used to evaluate association with toxicity. Genes are encoded as "yes" if phenotypic abnormalities in the following organs/systems are reported as significant: Liver, Spleen, Heart, Reproductive, Renal, Urinary, Skin, Neurological.

Small molecule druggability

Candidate Immunai targets are evaluated for targetability by a) cross-referencing candidate genes with tool compound availability and b) small molecule drugs by cross-referencing with curated functional annotations of genes (referencing GO and PANTHER functional annotations). DGIdb interaction database[11] (v. Dec2023) was used. 1411 druggable gene-products were selected based on availability of: Antibodies, Blockers, Inhibitors, Inverse agonists. GO and PANTHER functional annotations that were deemed to be druggable by small molecules.

Cell surface expression

Gene products are evaluated for targetability by presence of in Surfaceome database

SURFY Database. This is an *in silico* approach using domain-specific features. Surface definition manually curated by authors to refine entry for proteins with known localization. Database was curated to add "FPR_PNAS" cut off score as defined by Baush-Fluck, et al[12]. In addition, any gene that was functionally annotated with subcellular localization from HPA as "plasma membrane" was also included in the cell surface categorization. Subcellular location of each gene was further evaluated based on Human protein atlas[13] (v 23.0 Dec 2023).

Removal of housekeeping genes

Housekeeping genes were identified by cross-referencing those characterized by the GSEA human_housekeeping database. 7 genes were removed because of this (HLA-DRA,LGALS1,TXN,FYN,HLA-DQA1,LGALS3,CD3E)

Human intestinal protein expression

Expression of each gene was evaluated in intestinal tissues using the human protein atlas[13] (v 23.0 Dec 2023): Normal tissue data. Expression value is reported as the highest expression score in either rectum, colon, duodenum, or small intestine and protein levels with "Uncertain" Reliability are reported as NA. We scored each gene for qualitative intestinal expression (Not detected/Low/Medium/High).

#### Experimental functional validation protocols

CD14 monocyte differentiation and Endonuclease mediated knockout

Human PBMC from healthy volunteers were enriched for CD14+ monocytes by using the EasySepTM Human CD3 Positive Selection Kit II (to deplete CD3+ cells) followed by the EasySepTM Human Monocyte Isolation Kit (EasySepTM Human Monocyte Isolation Kit, STEMCELL, Technologies 19359). 6 donors were tested for each target. Nucleofection was performed using the P3 primary cell kit from Lonza (V4SP-3096), and sequence-specific ribonucleoprotein complexes targeting each locus (IDT, Table 6). Cells were then cultured in XVIVO media (TheraPEAK® X-VIVO® 15, LONZA BP04-744Q) + 1% human AB serum (Gemini Bio-Products, 100-512-100) + M-CSF (20ng/mL) for 5 days. On day 5, MNPs are polarized using either M-CSF (PeproTech AF-300-25, 20ng/mL) +IL-1β (PeproTech, AF-200-01B, 20ng/mL), +LPS (Invivogen, tlrl-3pelpsm, 5ng/mL) or GMCSF (PeproTech, AF-300-03, 20ng/mL) + IFN$\gamma$ (PeproTech, AF-300-02, 50ng/mL) +LPS (Invivogen, tlrl-3pelpsm, 5ng/mL). On day 7, the polarization media is refreshed, and cells are stimulated with the appropriate stimulus at the following concentrations: Iloprost (Sigma-Aldrich,SML1651, 0.1nM), IL27 (PeproTech, 200-38, 20ng/mL), IL6 (PeproTech, AF-200-06, 20ng/mL). On day 9, supernatants and cells are collected for readout.

Human fibroblast stimulation and Endonuclease mediated knockout

Human primary small intestinal fibroblasts (Cell Biologics H-6025, 2 donors) and, human primary colonic fibroblasts ( Cell Biologics H-6231, 2 donors) cells are expanded in Fibroblast Growth Medium (PromoCell Inc., C-23010) for 5 days. On experimental day 0, fibroblasts were nucleofected using the P3 primary cell kit from Lonza (V4SP-3096) and sequence-specific ribonucleoprotein complexes targeting each locus(IDT, Table 6) as described above. On day 6, the culture media is refreshed and IL6ST KO and F8 KO cells are stimulated with OSM (Peprotech, 300-10T, 100ng/mL) On day 9, supernatants and cells are collected for readout.

Antibodies and flow cytometry

Antibodies and reagents are listed in Table S7. In brief, cell suspensions were collected as described above, blocked with Fc Receptor Blocking Solution and stained with Viability dye in flow staining buffer (BD, 554656) . Cells are then stained with surface antibodies in flow staining buffer (BD, 554656) with 1:10 BV Buffer (BD 566385). For intracellular staining (ICS) of fibroblasts, cells were stained with surface markers, fixed permeabilized for 45 min at 4°C with Ebioscience™ Foxp3 / Transcription Factor Staining Buffer Set (Thermo Scientific, 00-5523-00). Cells were stained with ICS antibodies in permeabilization buffer for45 min at 4°C. Cells are resuspended in 100ul of FACS buffer with precision counting beads (BioLegend 424902). Flow cytometry was performed using a Cytek Aurora spectral flow cytometer, and data were analyzed with FlowJo. For each donor, cell type, target, and activation condition, the relative gMFI (geometric Mean Fluorescence Intensity) values of the markers were calculated using a background of the negative control gene (F8, B2M) knockout cells.

Secreted protein analysis, quality control and processing

Collected cell culture supernatant is centrifuged at 500rcf for 5min, collected and frozen at -80°C. Multiplexed cytokine assay was performed by Nomic Bio using their core panel of 275 analytes.

qPCR for target knockout efficiency

Cells are collected and lysed with RLT Buffer: RNeasy Lysis Buff (Qiagen 79216). Lysed cells are stored at -80°C until analysis. Total RNA was isolated using Qiagen RNeasy Kit (Cat no. 74106). For cDNA synthesis, a max of 1 μg total RNA was reverse-transcribed for 1 h at 37°C with an RNA-to-cDNA kit (Applied Biosystems, 4387406). For quantitative PCR, SYBR green qPCR master mix (New England Biolabs (NEB), M3003E ) and the following primers were used: human ACTIN forward, 5′-GATTCCTATGTGGGCGACGA-3′, and reverse, 5′-CACAGGACTCCATGCCCAG-3′; human PTGIR forward, 5′-:AGAGACACGAGGAGCAAAGC-3′, and reverse, 5′-CCACACCGGCCACGAACATC-3′, human IL6ST forward, 5′-GAACAGCATCCAGTGTCACC-3′, and reverse, 5′-AACACTCTTAATACTTGGGT-3′. To measure relative abundance of wild-type transcript versus edited transcript, we used custom designed primers. Typically, lead sgRNA sequence was used as the reverse primer and the forward primer sequence was determined using PrimerBlast (https://www.ncbi.nlm.nih.gov/tools/primer-blast/).

RNA library preparation for bulk-RNA sequencing

Total RNA was isolated from cultured cells using the Qiagen RNeasy® Plus 96 Kit (Qiagen, Cat. No. 74192) according to the manufacturer's protocol described in the RNeasy Plus 96 Handbook. Briefly, a minimum of 200,000 cells per sample were lysed in RLT Plus buffer (without β-mercaptoethanol), and genomic DNA was removed using the gDNA Eliminator 96-well plate. Lysates were processed through silica-membrane 96-well plates under vacuum or spin columns, and RNA was eluted in RNase-free water. RNA concentration was measured using the Qubit RNA HS Assay Kit (Thermo Fisher Scientific), and RNA integrity was evaluated using the Agilent TapeStation 4200 with the High Sensitivity RNA ScreenTape Assay (Agilent Technologies). Only samples with an RNA Integrity Number equivalent (RINe) ≥ 7.0 were used for library preparation. RNA-seq libraries were prepared using the Illumina Stranded mRNA Prep Kit (Illumina, Cat. No. 20040534) according to the manufacturer's instructions. Briefly, maximum 1000 ng of polyadenylated mRNA was enriched from total RNA using oligo(dT) magnetic beads, followed by fragmentation and first-strand cDNA synthesis using random priming. Second-strand synthesis was carried out to generate double-stranded cDNA incorporating dUTP to preserve strand specificity. The cDNA underwent end-repair, A-tailing, and ligation of unique dual-indexed adapters. Libraries were PCR-amplified and purified using KAPA HyperPure Beads (Roche, Cat. No. 08963860001). Final library quality and fragment size distribution were evaluated using the LabChip GX Touch HT Nucleic Acid Analyzer with the DNA High Sensitivity Assay (Revvity) or the Agilent TapeStation 4200 with the D5000 ScreenTape Assay (Agilent Technologies). Final library concentrations were determined by the Qubit 1X dsDNA High Sensitivity Assay Kit.

Final libraries were normalized and pooled equimolar for sequencing. Pooled libraries were sequenced on an Illumina NovaSeq System using a 25B flow cell with paired-end 100 bp or 150 bp reads.

Bulk RNA-seq data processing and analysis:

Illumina fastq files were processed to generate count matrices per experiment using our standardized bulk analysis workflow which includes read alignment, deduplication, mapping, and gene quantification. Genes with zero counts were excluded from downstream analysis. Per experiment, edgeR and limma were used to calculate paired differentially expressed genes between conditions (e.g., target versus control knockout) and log fold change between conditions per gene were calculated along with the nominal and adjusted p-values. Groups of gene programs were further tested for enrichment between conditions using fgsea and normalized enrichment scores and p-values were calculated per gene program. These were then visualized as heatmap or volcano plots to identify the least and most perturbed genes and gene sets per experiment.

<u>Clinical projection to the IBD tissue atlas and external validation cohort</u>

Gene signatures for the *in vitro* treatment or target KO RNA signatures are compiled from the differentially expressed genes. Upregulated and downregulated *in vitro*-derived signatures weighted by fold changes of individual genes were projected onto samples from the pseudobulked samples matched by cell type to the KO cell type in both the IBD atlas and external validation cohorts[14]. The enrichment of cell type specific IPR features-genes was tested for each cell type specific *in vitro* condition. The genes in each IPR feature were filtered for those with a correlation > 0.6, or a correlation > 0.3 and passing a MAD threshold. These filtered genes were defined as the IPR signature corresponding to each feature. The IPR signatures derived from each feature were used as an input to GSEA the differentially expressed genes in the Target KO vs control KO contrast.
